# Genetic diversity, structure, and kinship analysis of *Trachemys venusta venusta* in Wildlife Management Units and wild populations in south Mexico. Implications for conservation and management

**DOI:** 10.1101/2020.01.27.920983

**Authors:** Elsi B. Recino-Reyes, Julia M. Lesher-Gordillo, Salima Machkour-M’Rabet, Manuel I. Gallardo-Alvárez, Claudia E. Zenteno-Ruiz, León D. Olivera-Gómez, Alejandra Valdés-Marín, Guadalupe Gómez-Carrasco, Liliana Ríos-Rodas, María del Rosario Barragán-Vázquez, Raymundo Hernández Martínez

## Abstract

The Meso-American slider turtle (*Trachemys venusta*) is a freshwater turtle endemic to Mexico and Central America. Due to the overexploitation of its natural populations, it is in the at risk category formulated by the Official Mexican Standard NOM-059-ECOL-2010. In the state of Tabasco, Management Units for the Conservation of Wildlife (UMA) were created to reduce the impact of overexploitation of freshwater turtles. However, no genetic management plan was considered. This study presents the level of genetic diversity of the founder individuals in order to develop a management plan which will optimize reproduction in the UMA. Genetic diversity was compared between captive (n = 45) and wild (n = 86) individuals using 14 microsatellite molecular markers. Level of genetic diversity could be considered as low (*H*_e_ < 0.6) for a species of turtle and suggests that a higher level of protection is required for this particular species. Furthermore, values were slightly higher for the captive group reflecting the mix of genetic sources (founding individuals from different localities) and demonstrating that the captive population is genetically representative of natural populations. The genetic structure analysis revealed a relationship between captive and wild populations, indicating the influence of the two principal river basins in this region on the population of freshwater turtles. Finally, according to the results obtained from the analysis conducted using Storm and ML-Relate programs, we recommend the use of 19 females and 13 males, generating a potential of 247 dyads with no relationship. These first results of genetic management in a Mexican UMA, demonstrate the importance of molecular approaches at the time of managing and conserving species in captivity.

## Introduction

Currently, the world faces a higher and more rapid loss of its biodiversity [1], predominantly due to anthropogenic activities leading to climate change, habitat loss, introduction of invasive species, pollution, habitat degradation among others; consequently, captive breeding programs are a priority for species conservation [2]. During the last few decades, there has been an increase in global awareness on the need to conserve species through the management of captive populations and develop integrative conservation and management strategies [3–4]. Accordingly, the World Association for Zoos and Aquariums promotes the increase of animal management programs at population level to establish viable populations recognizing that this represents a biological and organizational challenge [3]. In addition, other initiatives are emerging such as the Community-Based Natural Resources Management that began in Africa and promoted the integration of the complex relationships among society (e.g., local livelihoods), economic and political authorities, environment, and sustainability principles for the management of natural resources [4]. In Mexico, the initiative of environmental policy started in 1997 with the creation of the National Program for Wildlife Conservation and Productive Diversification of the Rural Sector in which one of the axes is the development of the System of Management Units for Wildlife Conservation (SUMA, by its Spanish acronym) created by the Ministry of Environment and Natural Resources (SEMARNAT, by its Spanish acronym) [4–5]. Through this program, many Wildlife Management Units (UMAs, by its Spanish acronym) have been created [5–6]. The goal of UMAs is to provide an adequate and economically viable management of wildlife resources (fauna and flora) for conservation, rescue, and preservation purposes [7]. UMAs incorporate a wide range of activities such as research, recreation, environmental education, game farms, and commercialization of wildlife by products which are subject to regulated laws [6]. Furthermore, the creation of UMAs has favored the development of biological corridors where UMAs are interconnected with protected natural areas, as well as areas where sustainable use of the species is carried out, thus contributing to the improvement of biodiversity conservation [8]. Unfortunately, genetic aspects linked to captive breeding are rarely considered in the management of an UMA, which may lead to the failure of these programs.

One of the principal objectives of captive breeding programs is to obtain and maintain a population with a high level of genetic variation [9]. Generally, captive populations are small and established with few founders’ individuals, making these populations prone to high genetic damages such as a loss of genetic diversity, inbreeding depression, accumulation of novel deleterious mutations, and genetic adaptation to captivity [10]. However, from the few investigations conducted in UMAs, the genetic considerations (e.g., the genetic diversity of founders and relations of kinship) are one of the least considered aspects when establishing and managing an UMA. In Mexico, studies using genetic tools to improve the management of captive populations of UMA are scarce. Zarza et al. [11] identified the origin of 24 individuals of *Ctenosaura pectinata* Wiegmann 1834 (Squamata, Iguanidae), a threatened Mexican endemic spiny-tailed iguana, bred in two Mexican UMAs. The study was conducted using mtDNA and microsatellites molecular markers as well as genetic assignment methods, with the aim of comparing these individuals with a database of 341 individuals from 49 localities. This type of research demonstrates the importance of genetic tools in identifying the origin of individuals before a possible release in the wild.

Slider turtles of the genus *Trachemys* Agassiz, 1857 (Testudines, Emydidae) are freshwater turtles that present a wide distribution throughout the New World with 26 recognized extant forms [12]. In Mexico, a total of 13 subspecies are recognized, belonging to a variable number of species, from between one and nine, depending on the author concerned and partially dependent on the species concept used to identify lineages (review in [13]). One widely distributed *Trachemys* species, is *T. venusta* (Gray, 1856) with populations present from the Atlantic river basins of Mexico (and an isolated population in Acapulco) to Central America follow the most recent genetic analysis [13]. Its nomenclature is still a bit uncertain, therefore, genetic studies have been conducted in order to obtain a clearer understanding of its classification [14–16]. Three subspecies are described for *T. venusta* [17], all found in Mexico: *T. venusta venusta* (Gray, 1856), *T. venusta cataspila* (Günther, 1885), and *T. venusta grayi* which based on a genetic analysis, was recently proposed as a full species *T. grayi* (Bocourt, 1868) [15]. In the state of Tabasco only one subspecies was reported, *T. venusta venusta* [13, 17], commonly called *hicotea*.

The *hicotea* has been very important for the culture and gastronomy of the Maya people since the prehispanic period [18]. One of the few studies that deal with the exploitation of turtles in the state of Tabasco shows that the ingrained consumption of turtles persists despite the fact that the species is considered endangered [18]. Turtles are still in demand for commercial and consumption purposes. In addition to this culinary practice, urban growth has increased markedly since the mid-20^th^ century leading to high rates of deforestation [19–20]. The city of Villahermosa, the largest urban area in the state of Tabasco, is a perfect example of such environmental degradation; over a period of only 40 years, 4,008 ha of forest vegetation and 289 ha of wetlands have been lost [20]. The uncontrolled exploitation of freshwater turtles in Mexico, particularly in Tabasco, together with habitat loss, has resulted in many species of turtles being placed in the endangered category in the Official Mexican Standard (Norma Oficial Mexicana), the NOM-059-SEMARNAT-2010. This regulation establishes the protection of the environment and native species of wild flora and fauna of Mexico considering the following categories: endangered, threatened, under special protection, and probably extinct in the wild. *Trachemys venusta* (previously known as *T. scripta venusta* [21] is classified as “under special protection” [22].

Molecular tools allow us identify the origin of individuals when unknown (confiscated pets, illegal seizure) and to infer kin relationships among captive founder individuals [2]. For an adequate management of the breeding program, the knowledge of the relatedness between individuals is fundamental for success, because it allows a minimization of mean kinship and retains the maximum of genetic variation (review in [2]). Two different ways could be used to evaluate kin relationships: (1) the relatedness (*r*) which is a measure of the fraction of identical alleles shared by offspring, and (2) the relationship category which is a particular pedigree (genealogical) relationship such as full siblings or half siblings [23].

In this study, the genetic aspects of the founder individuals of *T. venusta venusta* from one UMA of the state of Tabasco were characterized and compared to a wild population, with the purpose of establishing an efficient management plan for reproduction. Ensuring mating between unrelated individuals and maintaining an adequate genetic variability in the individuals produced at the UMA will optimize a reintroduction program. Considering this, our particular objectives were (1) to determine the genetic diversity level of founder individuals of *Trachemys venusta venusta* in a UMA and compare it with wild individuals, (2) to evaluate intra- and inter- genetic structure of the UMA and wild population, (3) to determine kin relationships (relatedness and pedigree) between males and females in the UMA, and finally (4) our results will be discussed from a management and conservation perspective.

## Materials and methods

### Ethics Statement

Permission to collect specimens of *Trachemys venusta* was provided by the Mexican Ministry of Environment and Natural Resources (SEMARNAT). Experimental protocol was approved by Ethical Committee from the “Universidad Juarez Autonoma de Tabasco”, Mexico.

### Sampling sites and collection

The samples used for this study were obtained from one Wildlife Management Unit (UMA) and from three different wild localities (Fig 1). All samples were collected manually during 2017 and 2018 or using Fyke nets [24]. All wild turtles are between three to 10 years old according to their morphometric measures [25].

**Fig 1.**
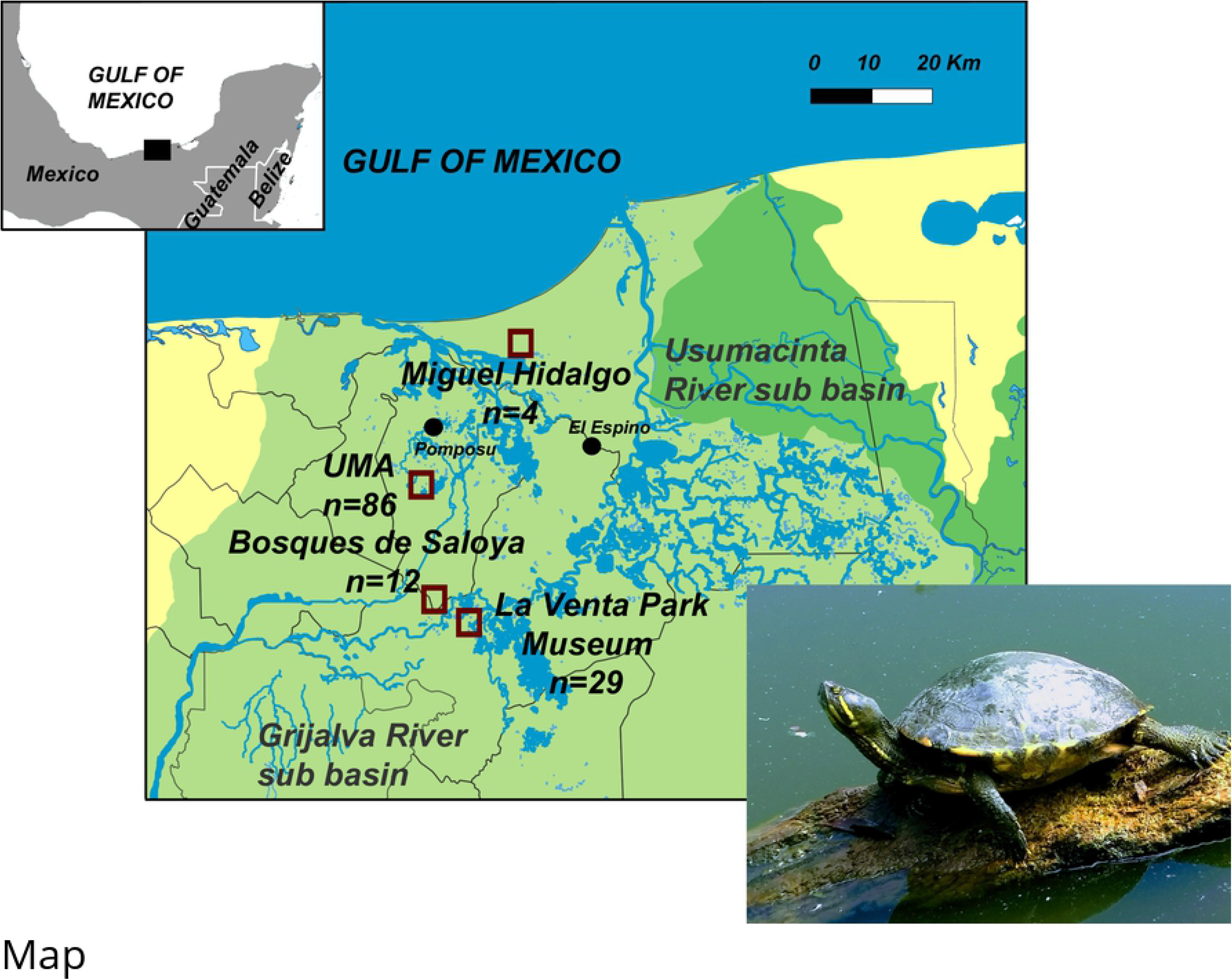
Study area in south Mexico. Squares represent location of the captive group (UMA) and the three localities for wild individuals (Miguel Hidalgo, Bosques de Saloya, and La Venta Park Museum) with their respective number of samples (n). The black points (Pomposu and El Espino) represent the localities of origin of founder’s individuals of captive groups (UMA) (photo by Julia M. Lesher-Gordillo).

The UMA is located in the state of Tabasco (municipality of Nacajuca; 18°11’23”N - 92°59’37”W) and it was created in 1978. It is dedicated to the reproduction of seven freshwater turtles *Dermatemys mawii* Gray, 1947 (Testudines, Dermatemydidae)*, Chelydra serpentine* L., 1758 (Testudines, Chelytridae)*, Staurotypus triporcatus* Wiegmann, 1828 (Testudines, Kinosternidae)*, Trachemys venusta venusta, Rhinoclemys aerolata* Duméril & Duméril, 1851 (Testudines, Geoemydidae)*, Claudius angustatus* Cope, 1865 (Testudines, Kinosternidae)*, Kinosternon leucostomum* Duméril & Duméril, 1851 (Testudines, Kinosternidae) for conservation purposes and to exchange individuals with other turtle farms. We obtained all initial founder individuals from the UMA (n = 86; n_female_ = 73 and n_male_ = 13). In 2008, this UMA reported 1650 newborn individuals of *T. venusta*, and the total number of individuals of this species was 4125 [26]. Furthermore, we obtained a total of 45 wild individuals (n_female_ = 32 and n_male_ = 13) from (1) La Venta Park Museum located in Tabasco state (municipality of Centro; 18°00’02”N - 92°56’08”W, n = 29), (2) Miguel Hidalgo (municipality of Centla; 18°20’00”N-92°30’00”W, n = 4), and (3) Bosques de Saloya (municipality of Nacajuca,18°11’22.85” N – 92°58’39.37” W, n = 12). Those individuals were considered as wild because they have been directly collected from a natural environment (e.g., Miguel Hidalgo and Bosques de Saloya), or because they have been recently rescued from the wild and donated directly to a center/farm (e.g., La Venta Park). Because some wild localities have a low number of samples, all the 45 wild individuals were pooled and analyzed as a single wild population.

Tissue samples were collected by small skin cuts (1 cm^2^) from the interdigital webbing. Before sampling, we disinfected the skin with 70% ethanol, subsequently we applied an antiseptic (methylthioninium chloride) on the wound to avoid infection. The collected samples were preserved in a tissue buffer solution (salt-saturated 20% DMSO; [27]), and stored at −80°C until DNA extraction.

### DNA extraction and microsatellite amplification

DNA was extracted using the DNeasy Blood and Tissue kit (QIAGEN); quality and concentration were verified using a UV-Vis spectrophotometer (Thermo Scientific NanoDrop One). We used a total of 14 microsatellite primers described as polymorphic for the genus *Trachemys* [28], one of them (TSC-108) was designed for this study using Primer3 software [29]. DNA amplification was performed at a final volume of 20μl per reaction, which contained: 20 ng of DNA, 2 μl of ultrapure water (INVITROGEN), 100 μM of forward primer, 100 μM of reverse primer, 15 μl of HotStarTaq® Master Mix Kit (QUIAGEN) (5 units/μl of polymerase, 20mM of TRIS-Cl, 100mM of KCl, 0.1mM EDTA and 400 μM of dNTPs). The amplifications were carried out in a Bio-Rad T100 thermocycler with the following conditions: 94°C for 7 min, followed by 45 cycles composed by denaturation at 92°C for 1 min, primer annealing temperature of 54°C or 55°C follow primers for 1 min, an extension at 72°C for 1 min, and a final extension at 72°C for 7 min. Horizontal electrophoresis was performed to visualize the PCR products using high definition agarose gels (agarose 1000, INVITROGEN) at a concentration of 3.4%. Gels were digitalized in a UV transilluminator Bio-print CX4 (VILBER LOUMAR) and it was interpreted using the BioVision software (VILBER LOUMAR). Microsatellites reproducibility was tested considering the allelic dropout (when a heterozygote is typed as a homozygote; [30]) and false allele (when homozygote is typed as a heterozygote; [30]) rates using GIMLET 1.3.3 [30]. The criteria used to test the genotyping error were: (1) analysis based on 5% of samples (eight individuals chosen at random) from the total sample size as suggested by Bonin et al. [31], (2) the same DNA extract was used in the reproducibility test and experiments, and (3) each DNA sample was replicated three times (twice in the same PCR run and once in the other PCR run) [32].

## Data analysis

### Genetic diversity

Genetic diversity was estimated considering two groups, UMA and wild population, through the number of alleles (*N_A_*), effective number of alleles (*N_e_*), observed heterozygosity (*H_o_*), and expected heterozygosity (*H_e_*) using Genalex 6.502 [33]. To test for differences in these parameters between the UMA and wild population, a Mann-Whitney *U* test was performed using Past 3.21 [34]. Because allelic richness (*N_A_*), as in other measures, does not consider the sample size, we determined the allelic richness (*AR*) following the rarefaction method implemented in HP-Rare 1.0 [35–36] which takes into account the difference in sample size. Additionally, the index of the polymorphic content (*PIC*) was determined using Cervus 3.0.7 [37]. Levels of inbreeding were estimated using three different parameters: at the population level we determined (1) the inbreeding coefficient (*F*) using Genalex 6.502, at the individual level we determined (2) the internal relatedness (*IR*) which evaluates the relatedness of the parents of an individual [38] (outbred individuals present values equal to or below zero, and positive values indicate some relatedness of parents with a maximum value of 1), and (3) the homozygosity by loci (*HL*) which is a homozygosity index that weights the contribution of each locus depending on their allelic variability [39], their values range from 0 (all loci are heterozygous) to 1 (all loci are homozygous). The Hardy-Weinberg equilibrium (*HWE*) for heterozygote deficit was tested with Genepop 4.0.10 [40–41], and the presence of null alleles at each loci was estimated with FreeNa [42]. Finally, we evaluated the effective population size (*N_E_*) using NeEstimator 2.1 [43] considering the linkage disequilibrium method with the lowest allele frequency at a critical level of 0.05, and the jackknife method to evaluate the 95% of confident interval (CIs) as recommended [44].

To test a possible bottleneck event, we compared the levels of *H_e_* excess related to the expected equilibrium heterozygosity (*H*_eq_) using Bottleneck 1.2.02 [45–46]. Two models of mutational equilibrium were considered: a mutational stage model (SMM) and a two-phase model (TPM), with 95% of the mutation assumed in a single step for the TPM model and a variance among multiple steps of 12 as recommended [46]. Significance was evaluated from the one-tailed Wilcoxon rank test which is more appropriate and powerful for less than 20 microsatellites [46]. Bottleneck event will be considered only if both models (SMM and TPM) are significant.

### Genetic structure

To determine the level of genetic differentiation between the UMA and the wild population, the values of *F_ST_* was estimated with Genalex 6.502. To identify the genetic structure between the UMA and the wild population, the Bayesian analysis implemented in Structure 2.3.1 [47] was applied. This method allows to determine the optimal number of groups/clusters (*K*) and assign to each individual a membership probability (*q_i_*) to new clusters. The admixture and allele correlated frequency models were used. To determine the optimal number of clusters, the program was executed ten times for different numbers of *K* (*K* from 1 to 5), and for each run the Markov Chain Monte Carlo (MCMC) algorithm was executed with a burn-in period of 100,000 steps followed by 100,000 steps. We used the Evanno method (Δ*K* method; [48]), implemented in the Structure Harvester website [49] to determine the best value of *K* that fit with our data. Because this method assigns individuals by force to the number of clusters considered at optimal, we decided to conduct a principal coordinate analysis (PCoA) to investigate genetic structure without a priori assignment. Finally, a hierarchical analysis of molecular variance (AMOVA; 9,999 permutations) was processed considering region level (UMA *vs* wild populations) and population level (different wild localities) using allelic distance matrix as input. Those ultimate analyses (PCoA and AMOVA) were conducted using Genalex 6.502.

### Kinship analysis

To evaluate pedigree relationships in the UMA, we used the ML-Relate program [37] which calculates the likelihood of each kind of relationship for each pair of individuals considering the following pedigree relationships: unrelated (U), half-siblings (HS), full-siblings (FS), and parents-offspring (PO). For each pair of individuals, the program provides by default, the highest likelihood, then suggests the highest relationship. ML-Relate provides a list of several possible relationships (the putative corresponding to the highest likelihood, and the alternatives) for each individual pair based on a test (statistical test and simulations) that determined which relations are consistent with the data (confidence set option; 0.05 level of significance and 1,000 simulations) [37]. Based on those relationships for each female-male pair from the UMA, we used a likelihood ratio test (1,000 simulations; specific hypotheses test option) to obtain a *P* value: if *P* is small (*P* < 0.05) the alternative hypothesis is rejected, thus we maintained the highest relationship proposed by the highest likelihood, and if *P* value is large (*P* > 0.05), putative and alternative relationships are consistent with the data, therefore, we selected the lowest relationship (U < HS < FS < PO) [37].

Furthermore, relatedness coefficient (*r*) among each female-male pair in the UMA was evaluated using Storm [50] and ML-Relate programs.

## Results

### Genetic diversity

A total of 86 individuals from the UMA and 45 from the wild population were successfully amplified with 14 microsatellite loci. However, considering that four loci were identified as possible null alleles (Table 1), we decided to eliminate them and make the following analysis only considering the 10 loci without null alleles. Genotyping error rates could be considered null (average allelic dropout rate of 0% and average false allele rate of 0%; because the errors were not significant) when compared with other studies (see Table 4 in [51]).

**Table 1.**
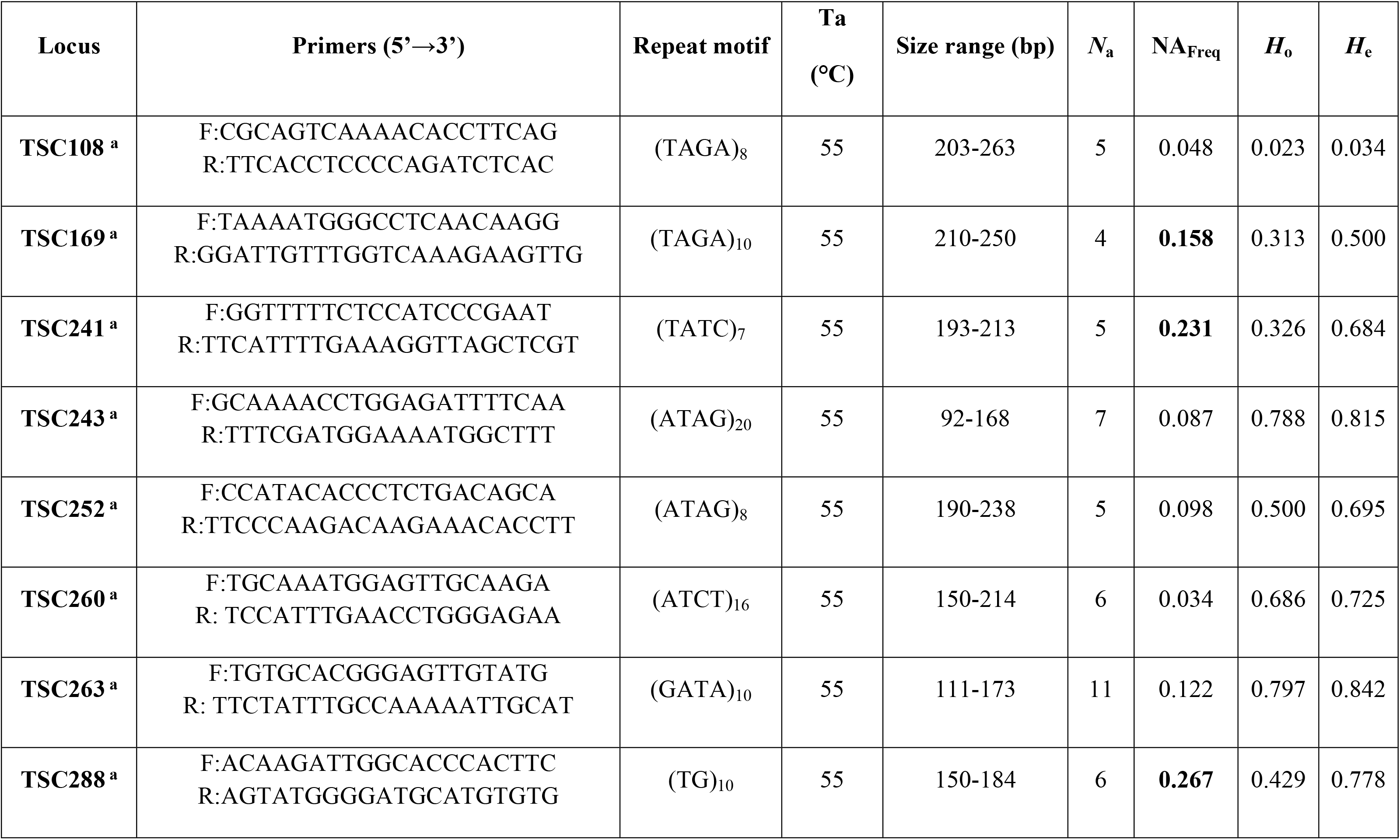
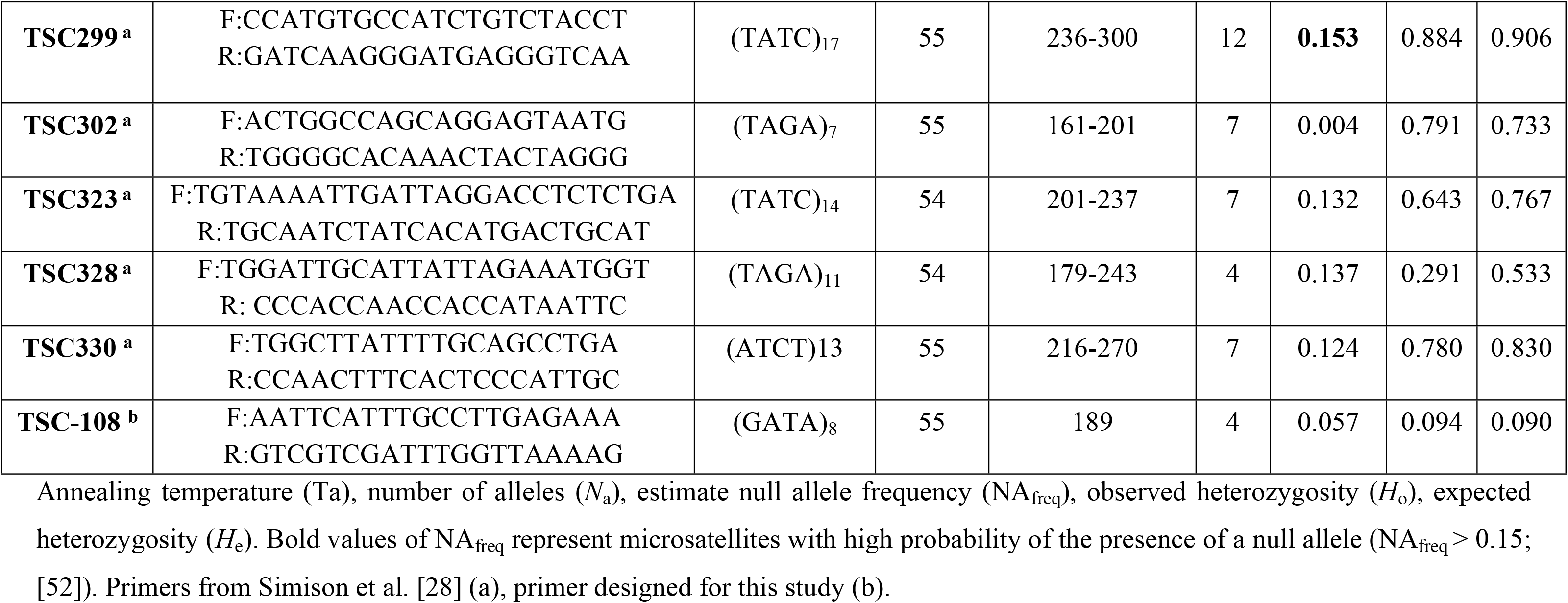
Characteristics of the 14 microsatellites used for *Trachemys venusta venusta* in south Mexico over the whole dataset.

Globally, genetic diversity parameters (*N*_a_, *N_e_*, *H*_o_, *H*_e_, *PIC*) were slightly higher for the UMA than for the wild population but never significant (Table 2). The allelic richness (*AR*) was very similar between both groups (UMA and wild populations), and this value that considers sample size does not differ from the value for the number of alleles (Table 2). The effective population size (*N*_E_) was low, with a value of 91 for the UMA and 110 for wild populations. The inbreeding coefficients (*F*) were positive for both groups but with a lower value for the UMA than for the wild population, and the HWE test for deficiency of heterozygotes was only highly significant for the wild population (Table 2). At the individual level, internal relatedness (*IR*) was very low for the UMA and high for the wild population with high individual variability for both groups (S1 Fig). The level of homozygosity (*HL*) was high for the UMA (an average of 33% of loci are homozygous per individual) and very high for the wild population (an average of 55% of loci are homozygous per individual) (Table 2), and even if some individual variability could be observed (S2 Fig) wild individuals clearly show a tendency to a high value of *HL*. No recent bottleneck was detected for any population as suggested by the L-shape graph (Fig 2) that shows a characteristic L-shape distribution (alleles with low frequency are the most numerous), and by both heterozygote excess tests (SMM and TPM) (Table 2).

**Fig 2.**
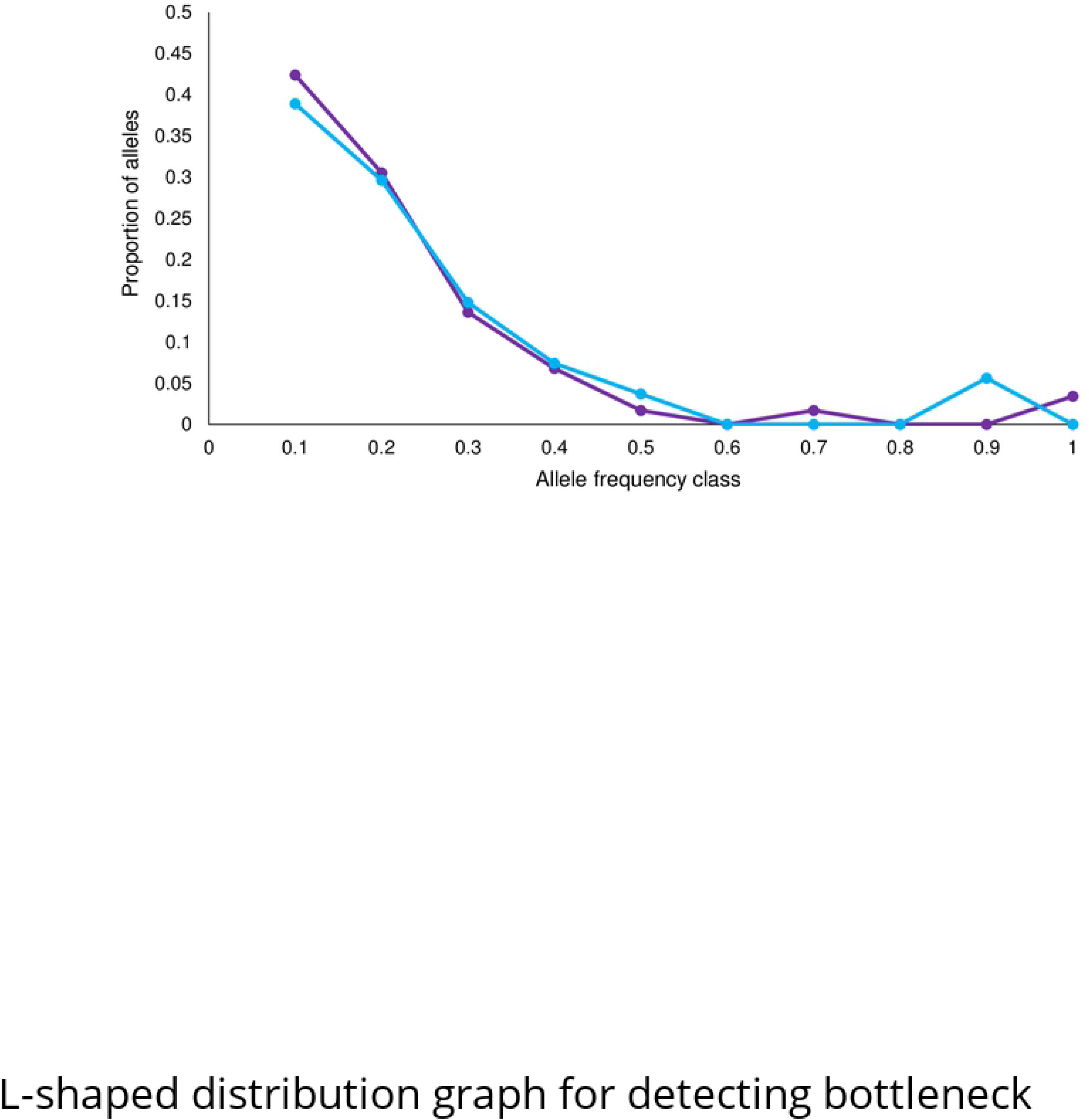
L-shaped distribution graph for detecting bottleneck considering the ten microsatellites for *Trachemys venusta venusta*. UMA (purple) and wild populations (blue).

**Table 2.**
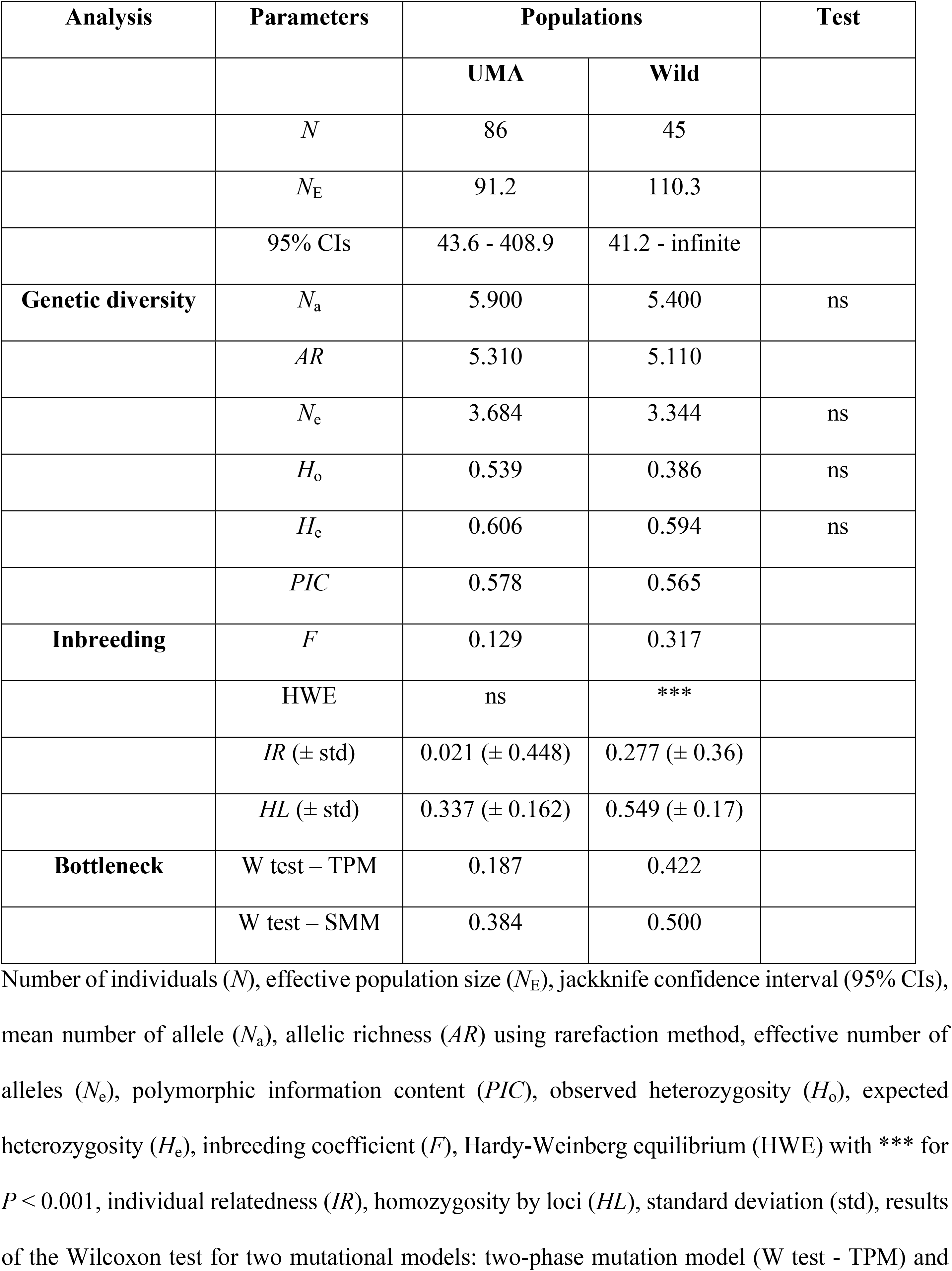
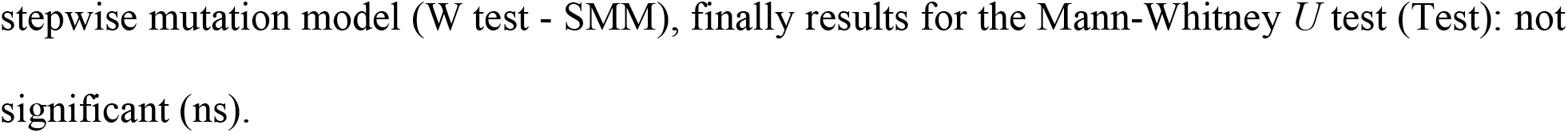
Statistical summary for *Trachemys venusta venusta* in south Mexico using ten polymorphic microsatellites for founder individuals of the UMA and for wild population.

| Analysis | Parameters | Populations |  | Test |
| --- | --- | --- | --- | --- |
|  |  | UMA | Wild |  |
| | $N$ | 86 | 45 | |
| | $N_E$ | 91.2 | 110.3 | |
|  | 95% CIs | 43.6 - 408.9 | 41.2 - infinite |  |
| **Genetic diversity** | $N_a$ | 5.900 | 5.400 | ns |
| | $AR$ | 5.310 | 5.110 | |
| | $N_e$ | 3.684 | 3.344 | ns |
| | $H_o$ | 0.539 | 0.386 | ns |
| | $H_e$ | 0.606 | 0.594 | ns |
| | $PIC$ | 0.578 | 0.565 | |
| **Inbreeding** | $F$ | 0.129 | 0.317 | |
|  | HWE | ns | *** |  |
| | $IR (\pm \text{std})$ | 0.021 ($\pm 0.448$) | 0.277 ($\pm 0.36$) | |
| | $HL (\pm \text{std})$ | 0.337 ($\pm 0.162$) | 0.549 ($\pm 0.17$) | |
| **Bottleneck** | W test – TPM | 0.187 | 0.422 |  |
|  | W test – SMM | 0.384 | 0.500 |  |

Number of individuals ( $N$ ), effective population size ( $N_E$ ), jackknife confidence interval (95% CIs), mean number of allele ( $N_a$ ), allelic richness ( $AR$ ) using rarefaction method, effective number of alleles ( $N_e$ ), polymorphic information content ( $PIC$ ), observed heterozygosity ( $H_o$ ), expected heterozygosity ( $H_e$ ), inbreeding coefficient ( $F$ ), Hardy-Weinberg equilibrium (HWE) with \*\*\* for $P < 0.001$ , individual relatedness ( $IR$ ), homozygosity by loci ( $HL$ ), standard deviation (std), results of the Wilcoxon test for two mutational models: two-phase mutation model (W test - TPM) and
stepwise mutation model (W test - SMM), finally results for the Mann-Whitney $U$ test (Test): not significant (ns).

### Genetic structure

The *F*_ST_ value between UMA and wild population was low (0.031) but significant (*P* = 0.0001), just as the results of AMOVA at the region level presented only 2% of variance (*P* = 0.0017) and 4% for populations level (*P* = 0.001), the rest of the variance corresponded to the individual variance (see details of analysis in S1 Table). The Bayesian analysis performed by the Structure program identified an optimal number of groups at *K* = 2 using the Δ*K* method (S3 Fig). All individuals were relatively well partitioned into the two new clusters according to their *Q* probabilities (Table 3), showing a good separation between individuals from the UMA group (purple in Fig 3) and individuals from the wild population (blue in Fig 3). Nevertheless, some individuals of the UMA present a genetic profile characteristic of a wild population. A subsequent Structure analysis for each new cluster did not permit the identification of a sub-structure (not shown). As observed for Bayesian analysis, PCoA analysis showed a relatively good separation between UMA (purple in Fig 4) and wild individuals (blue in Fig 4). Individuals from the La Venta locality (blue triangle) are more separated than others (blue circle and square).

**Fig 3.**
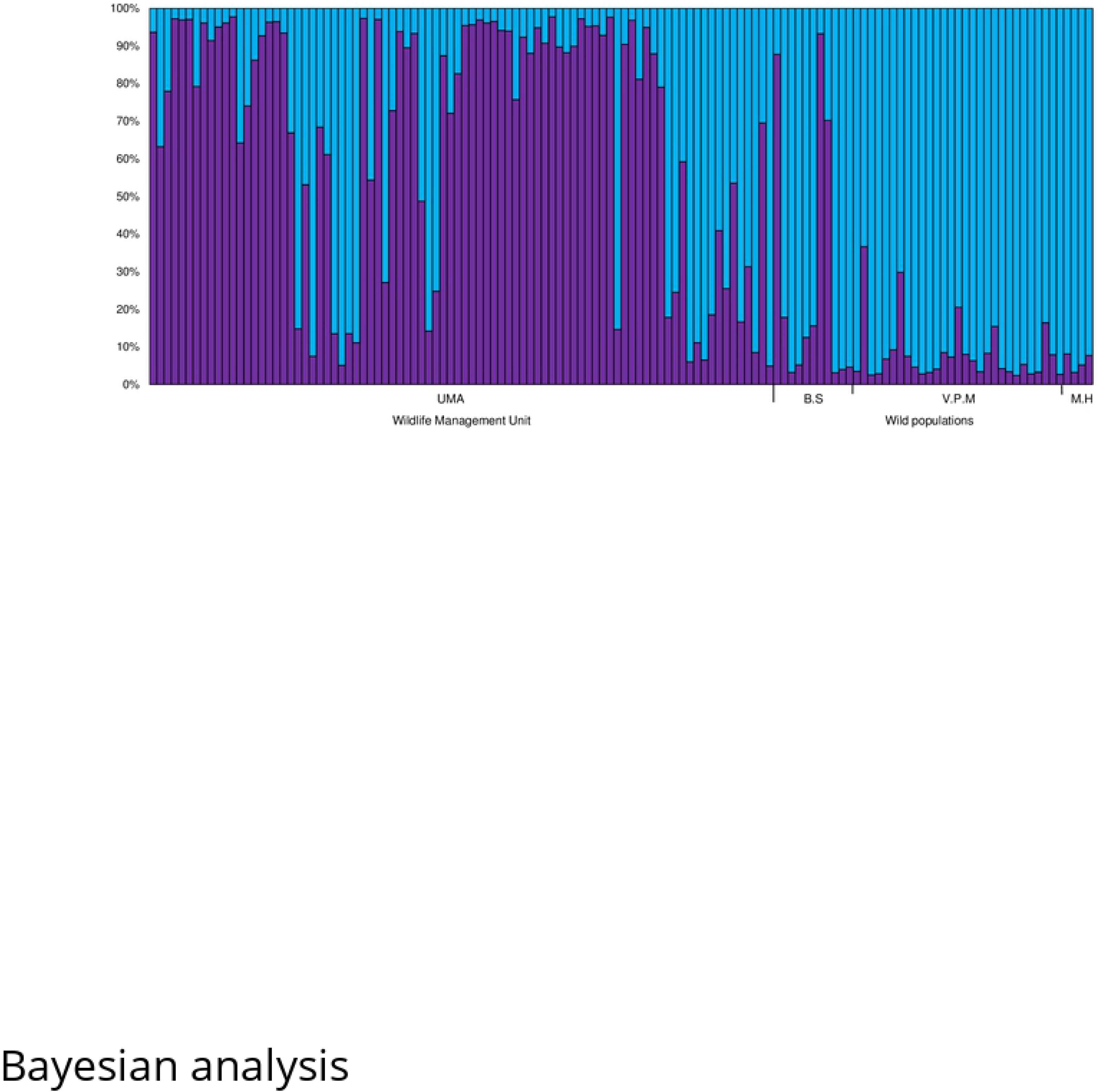
Bayesian analysis computed by Structure 2.3.3 software using *K* = 2 for *Trachemys venusta venusta*. Each individual is represented by a vertical line fragmented into k segments of length proportional to the estimated membership probability (*q*_i_) in the two new clusters. Cluster 1 characteristic of captive group (UMA; purple) and cluster 2 characteristic of wild individuals (bleu) with details of different localities: Bosques de Saloya (B.S), Venta Park Museum (V.P.M), and Miguel Hidalgo (M.H).

**Fig 4.**
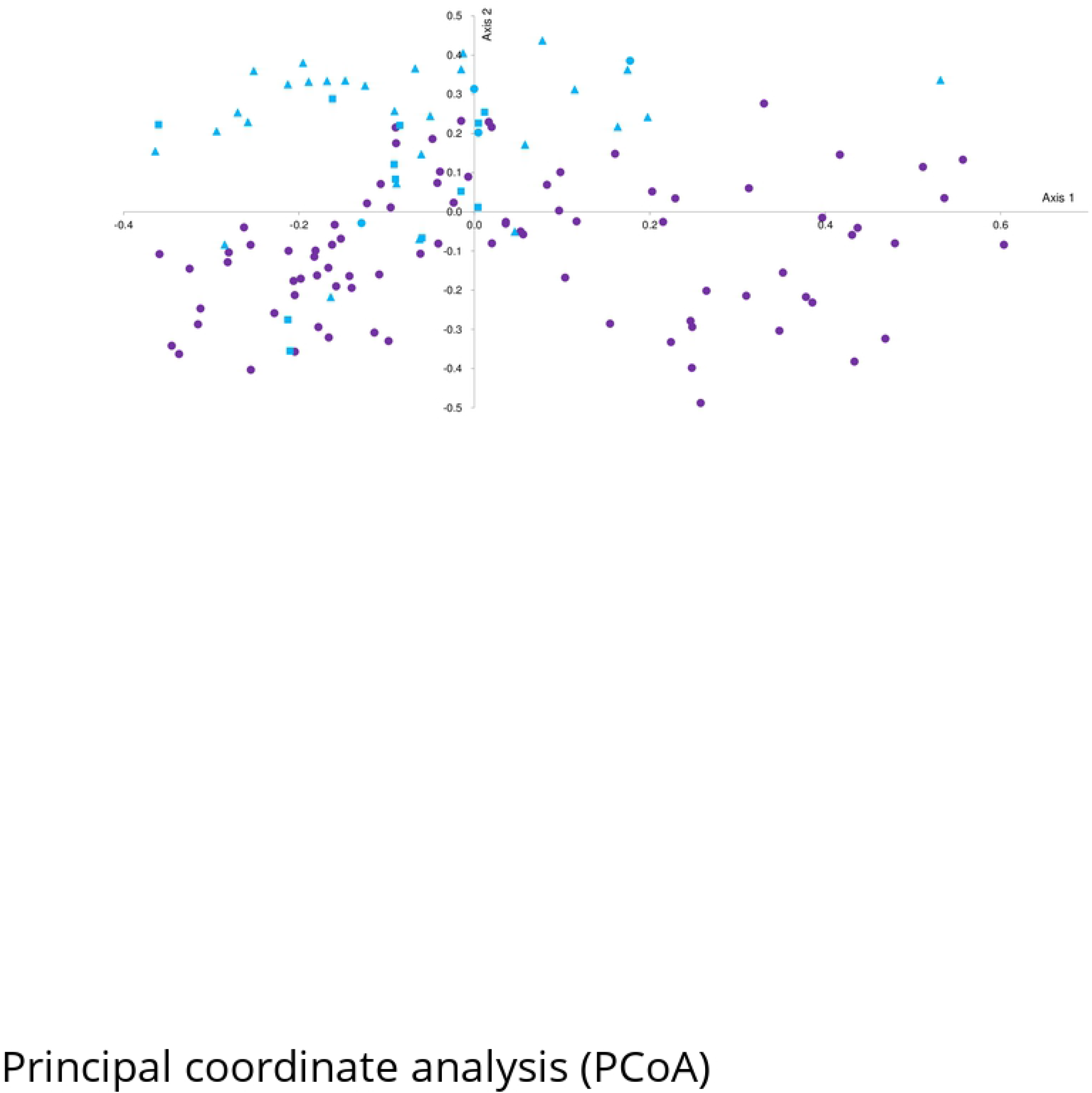
Principal coordinate analysis (PCoA) for *Trachemys venusta venusta* based on ten microsatellites. Individuals from Wildlife Management Unit (UMA; purple circle), and wild individuals (blue), localities: La Venta (triangle), Miguel Hidalgo (circle), and Bosques de Saloya (square).

**Table 3.** Probability *Q* of membership of individuals in each initial group to each of the inferred clusters determined by Structure 2.3.1 program.

| Initial group | Inferred clusters |  |
| --- | --- | --- |
|  | 1 | 2 |
| **UMA** | **0.680** | 0.320 |
| **Wild population** | 0.139 | **0.861** |

The dominant percentage of a group that is assigned to one cluster is indicated in bold.

### Kinship analysis in UMA

Relatedness coefficient (*r*) determined using Storm and ML-Relate clearly shows some kinship relations in the initial founders of the UMA. Storm and ML-Relate analysis presents values from −4.78 to 0.67 and from 0 to 0.63 respectively, considering all kind of dyads (female x female, male x male, and female x male). However, considering that we are interested in genetic management to optimize reproduction, we focused our analysis on female x male pairs only, which represent a total of 949 dyads. Using the relatedness coefficient generated by Storm we determined the proportion of each pedigree relation (PO, FS, HS, and U) based on the follow criteria: values ≤ 0 are assimilated to unrelated (U), values ≤ 0.25 are assimilated to half-sibling (HS), and values > 0.25 are assimilated to FS or PO without distinction [53–54]. We obtained 45% of unrelated (n = 427), 39% of HS (n = 367), and 16% of FS or PO (n = 155). Those pedigree predictions differed considerably from ML-Relate relationships, based on the highest likelihood which give us 79.7% of U, 15.4% of HS, 3.8% of FS, and 1.2% of PO for the same dataset (a total of 193 pairs of individuals presented half-sibling or greater kinship relation). Relatedness coefficients corresponding to those pedigree predictions were highly variable (Fig 5, full color histogram) with a mean of 0.039 (± 0.067SD, min = 0.000, max = 0.474) for U (blue in Fig 5), 0.230 (± 0.068SD, min = 0.087, max = 0.388) for HS (yellow in Fig 5), 0.383 (± 0.083SD, min = 0.230, max = 0.625) for FS (green in Fig 5), and 0.536 (± 0.046SD, min = 0.500, max = 0.611) for PO (red in Fig 5). When the likelihood ratio test was applied to compare the putative relationship (highest value of likelihood) with the alternative ones proposed by the confidence interval, we obtained: 91.0% of U, 7.5% of HS, 1.4% of FS, and 0.1% of PO for the same dataset (a total of 85 pairs of individuals presented half-sibling or greater kinship relation), also with a highly variable relatedness coefficient (Fig 5, hatched color histogram) with a mean of 0.060 (± 0.085SD, min = 0.000, max = 0.500) for U (blue in Fig 5), 0.342 (± 0.067SD, min = 0.228, max = 0.511) for HS (yellow in Fig 5), 0.509 (± 0.099SD, min = 0.340, max = 0.625) for FS (green in Fig 5), and for PO only one data (red in Fig 5).

**Fig 5.**
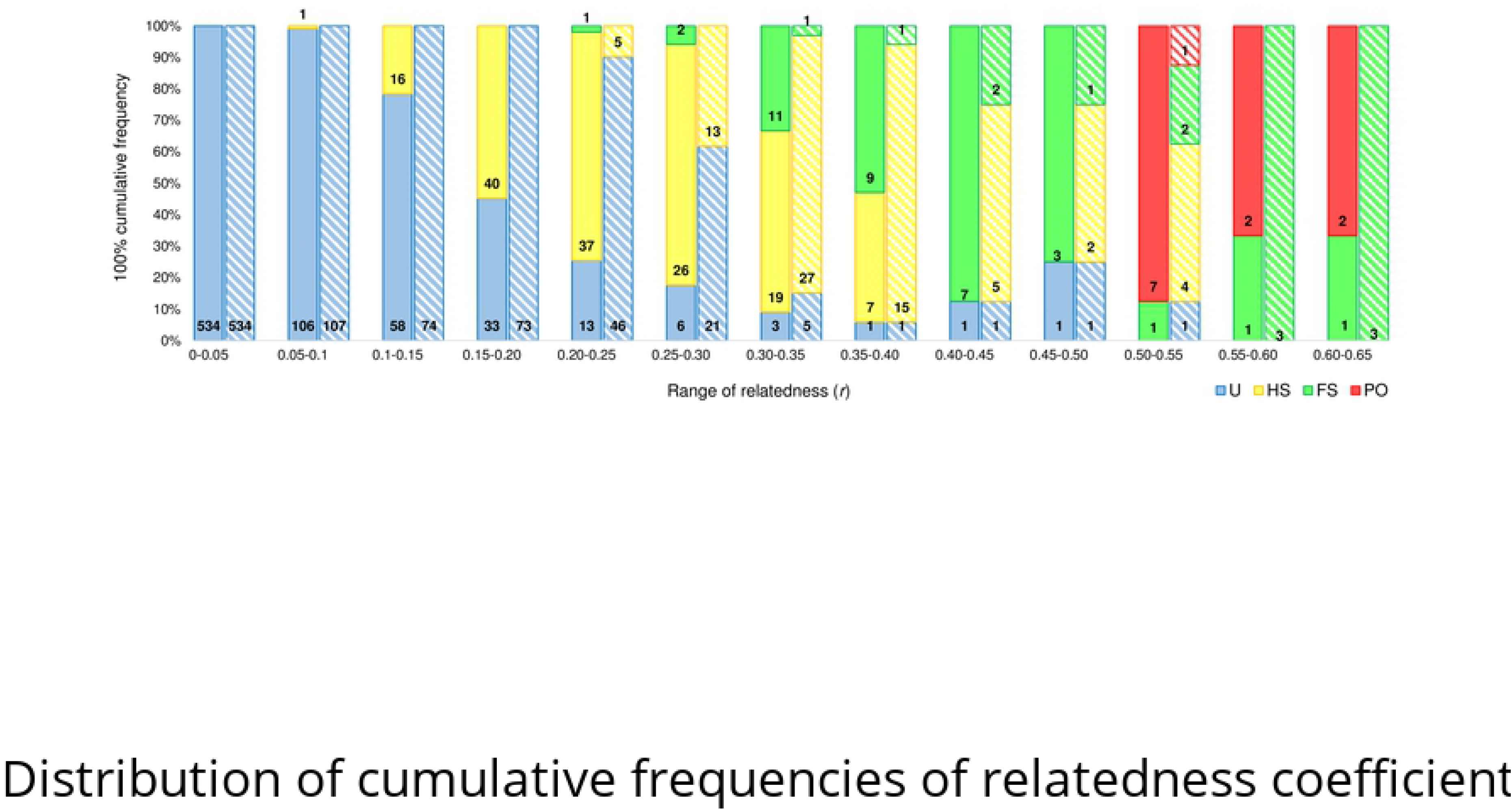
Distribution of cumulative frequencies of relatedness coefficient (*r*) determined by ML-RelateD for female x male dyads of *Trachemys venusta venusta* in the UMA. Estimation based on the highest likelihood (full color) and the estimation obtained after the likelihood ratio test (hatched color). Four relationships were considered: unrelated (blue), half-sibling (yellow), full-sibling (green), and parent-offspring (red). Numbers inside each bar of the histogram correspond to the number of pairs observed in this category.

To optimize reproduction with the best females and males in terms of kinship relation, we analyzed all female x male dyads (total of 949) and identified which pairs presented half-sibling or greater kinship relation. In the first instance, we considered kinship relation based on the highest likelihood prediction (letters of kinship relation type in bold in Table 4) and subsequently, we looked at which of these relationship could be changed to unrelated considering the likelihood ratio test results (letters of kinship relation type in grey in Table 4). From 73 female founders of the UMA, only 19 have no strict kin relationship with the 13 males (females highlighted in grey in Table 4). All other females present a minimum of one kin relationship with the male founders of the UMA.

**Table 4.**
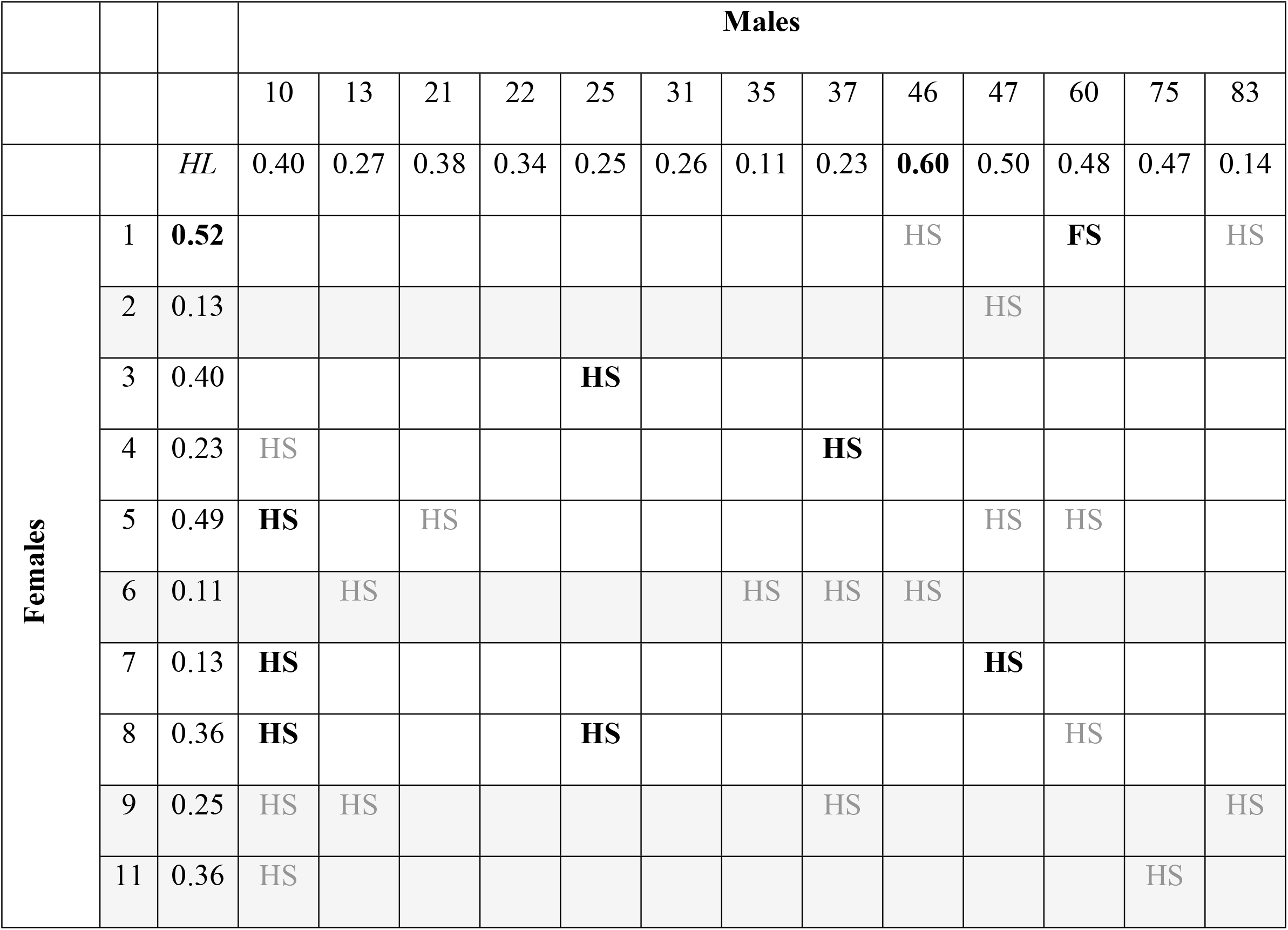
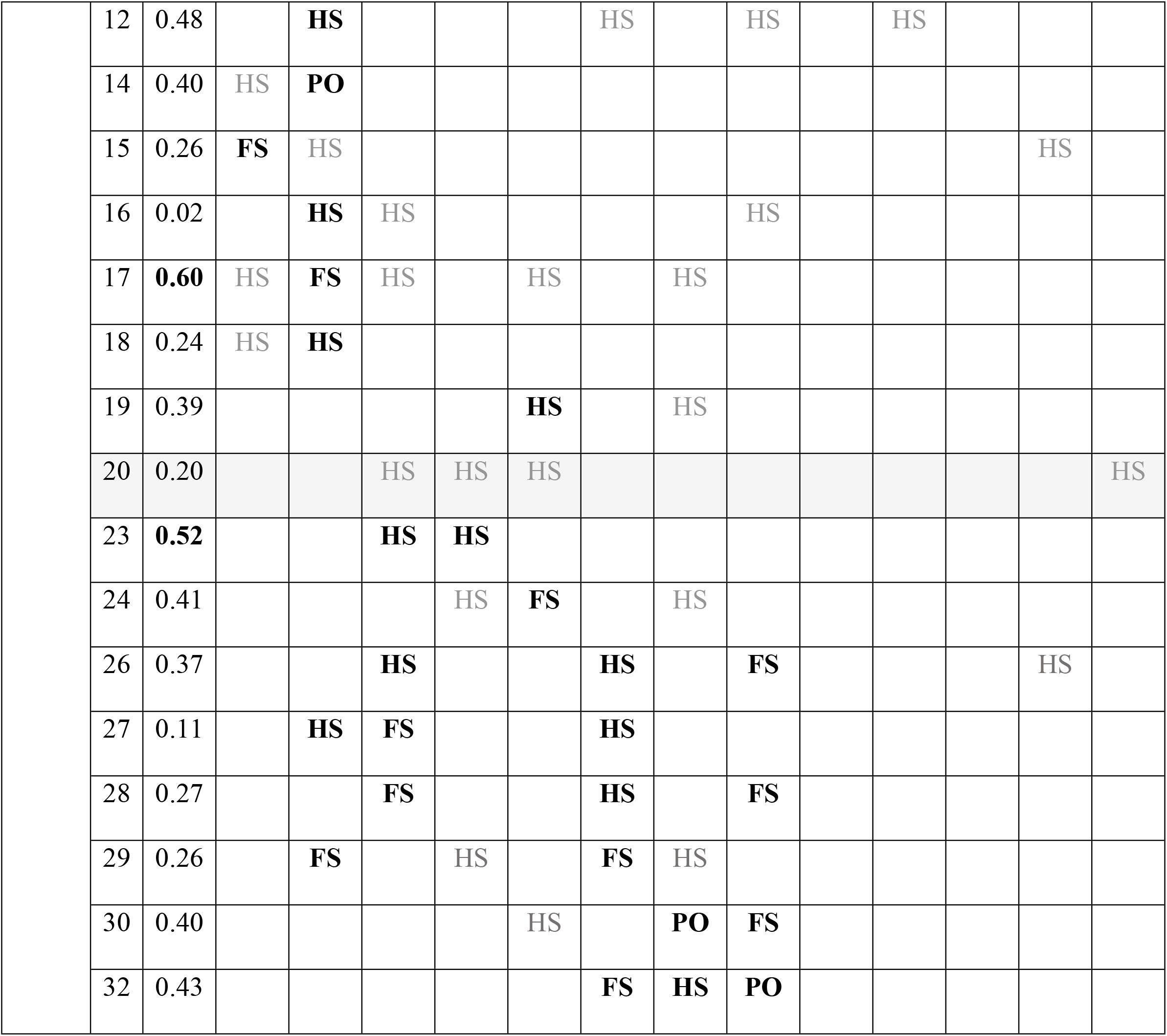
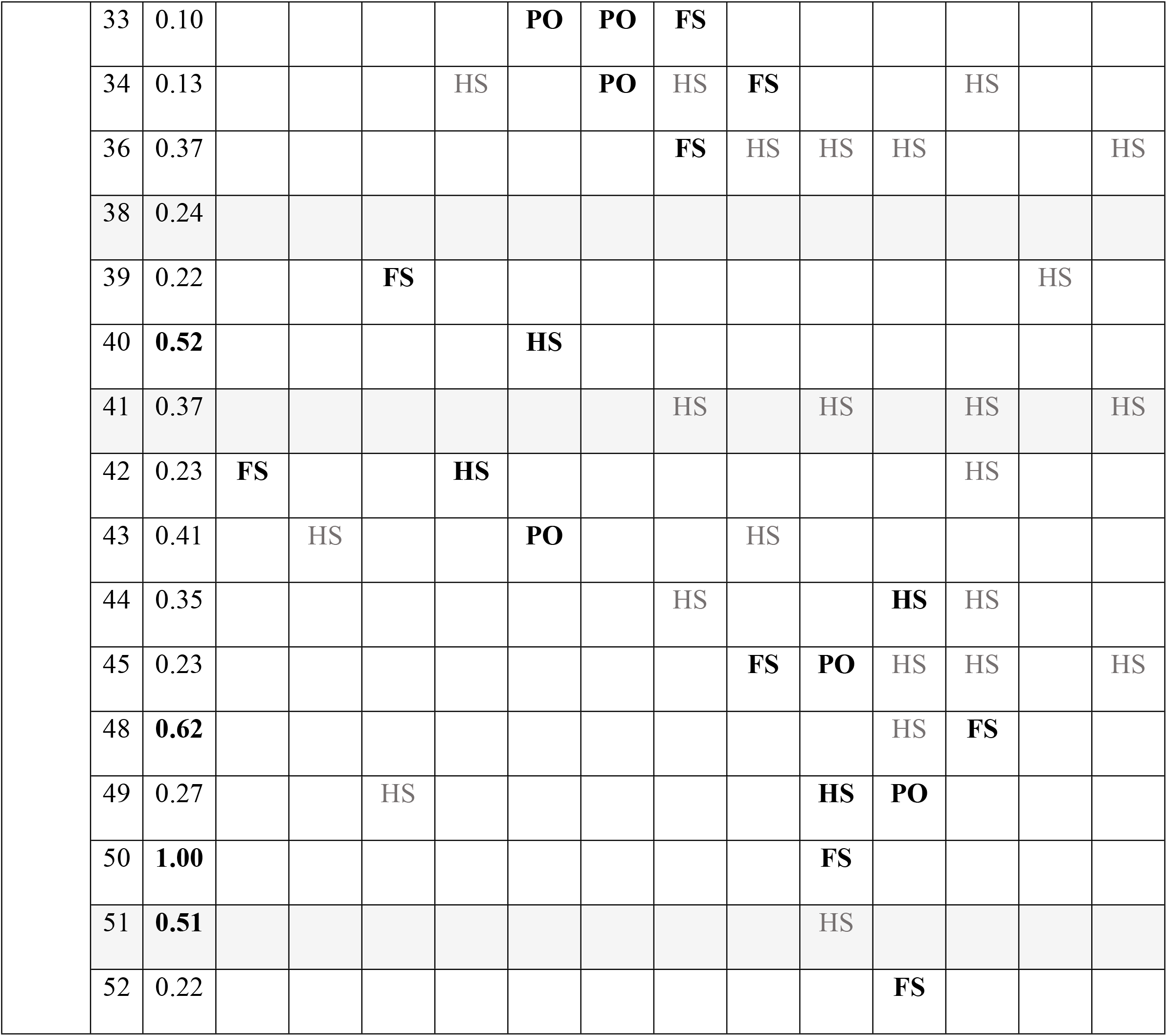
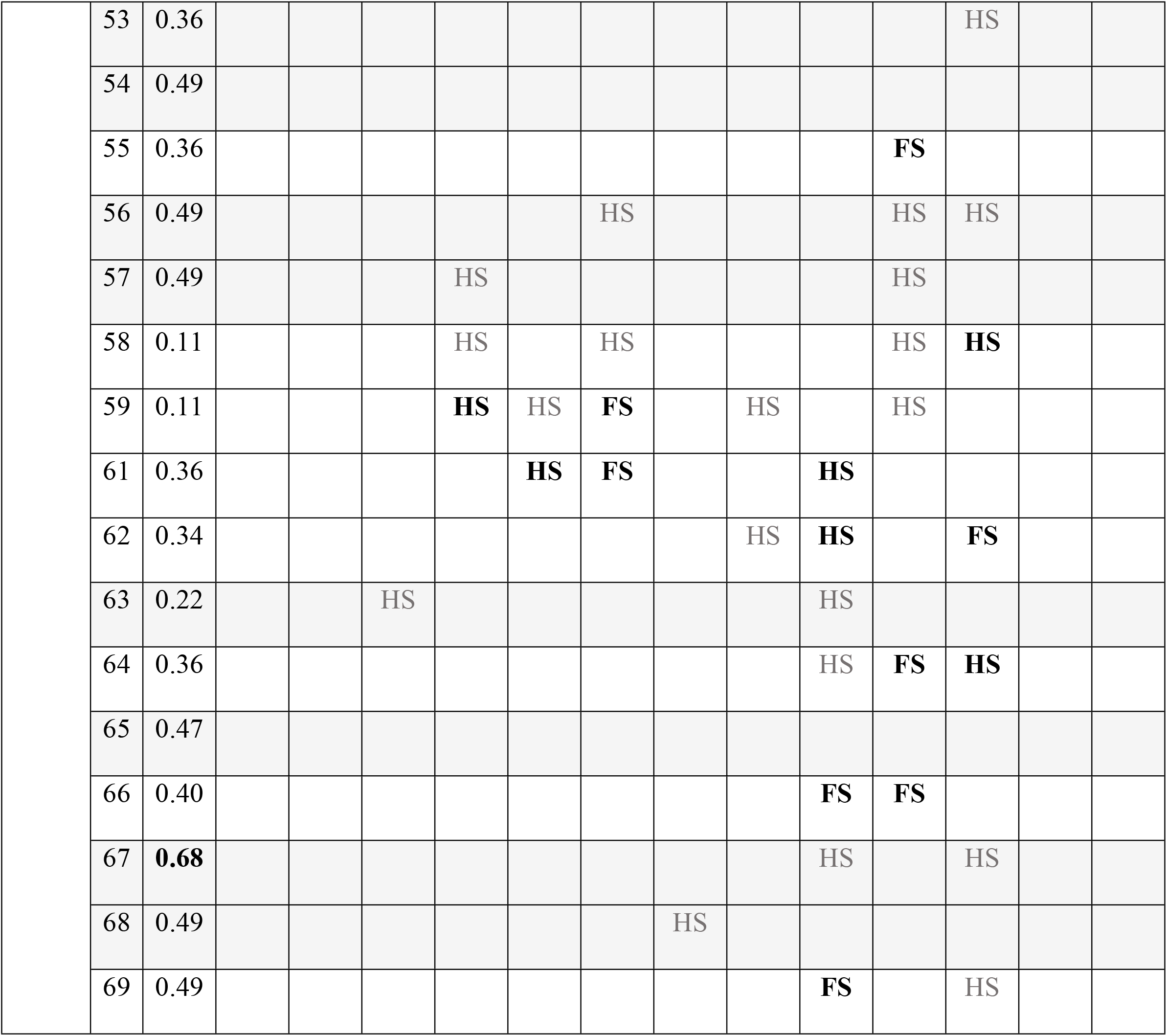
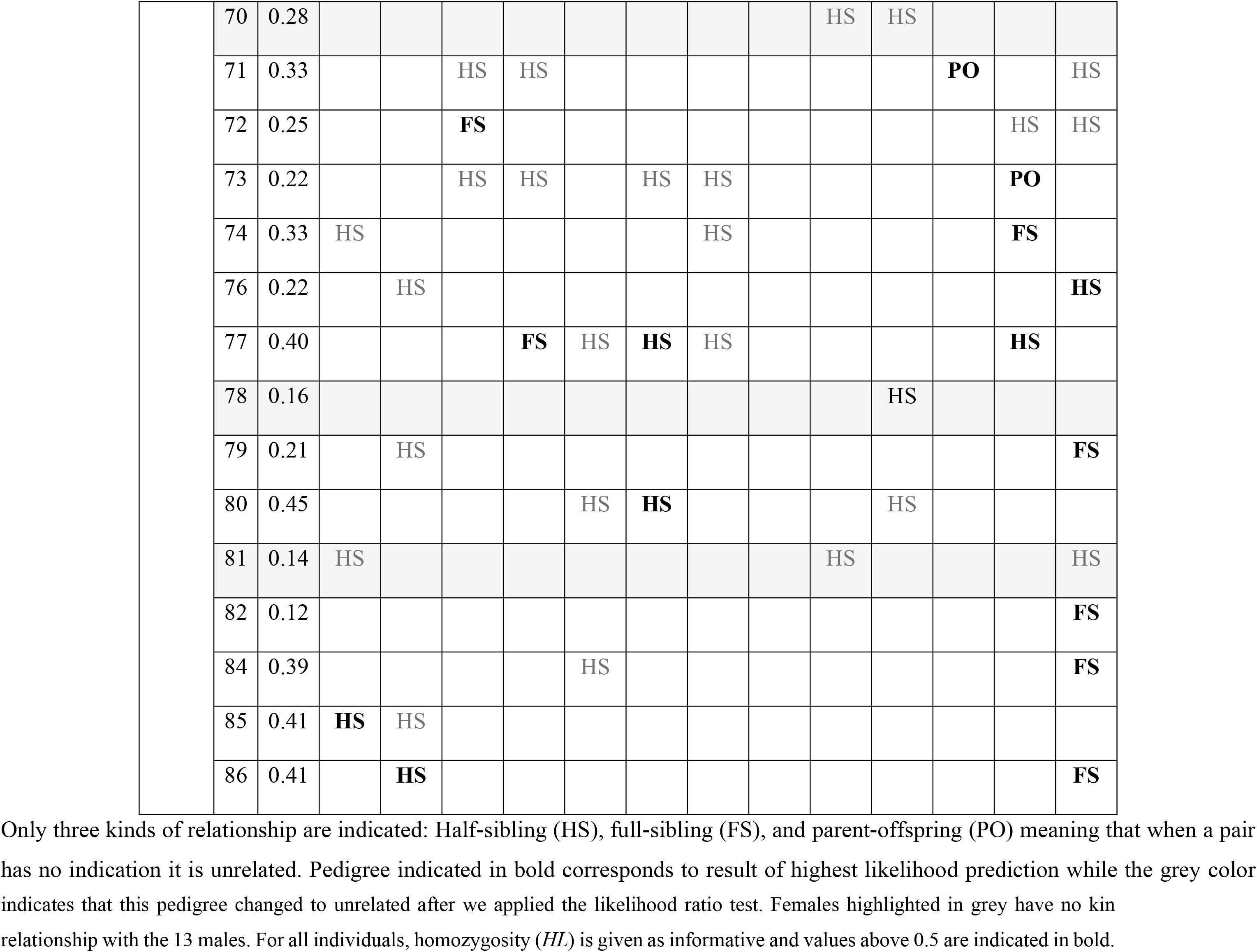
Pedigree relationships for all female x male dyads (n = 949) of *Trachemys venusta venusta* of the UMA determined by ML-Relate.

|  |  |  | Males |  |  |  |  |  |  |  |  |  |  |  |  |
| --- | --- | --- | --- | --- | --- | --- | --- | --- | --- | --- | --- | --- | --- | --- | --- |
|  |  |  | 10 | 13 | 21 | 22 | 25 | 31 | 35 | 37 | 46 | 47 | 60 | 75 | 83 |
|  |  | *HL* | 0.40 | 0.27 | 0.38 | 0.34 | 0.25 | 0.26 | 0.11 | 0.23 | **0.60** | 0.50 | 0.48 | 0.47 | 0.14 |
| Females | 1 | **0.52** |  |  |  |  |  |  |  |  | HS |  | **FS** |  | HS |
|  | 2 | 0.13 |  |  |  |  |  |  |  |  |  | HS |  |  |  |
|  | 3 | 0.40 |  |  |  |  | **HS** |  |  |  |  |  |  |  |  |
|  | 4 | 0.23 | HS |  |  |  |  |  |  | **HS** |  |  |  |  |  |
|  | 5 | 0.49 | **HS** |  | HS |  |  |  |  |  |  | HS | HS |  |  |
|  | 6 | 0.11 |  | HS |  |  |  |  | HS | HS | HS |  |  |  |  |
|  | 7 | 0.13 | **HS** |  |  |  |  |  |  |  |  | **HS** |  |  |  |
|  | 8 | 0.36 | **HS** |  |  |  | **HS** |  |  |  |  |  | HS |  |  |
|  | 9 | 0.25 | HS | HS |  |  |  |  |  | HS |  |  |  |  | HS |
|  | 11 | 0.36 | HS |  |  |  |  |  |  |  |  |  |  | HS |  |
|  | 12 | 0.48 |  | **HS** |  |  |  | HS |  | HS |  | HS |  |  |  |
|  | 14 | 0.40 | HS | **PO** |  |  |  |  |  |  |  |  |  |  |  |
|  | 15 | 0.26 | **FS** | HS |  |  |  |  |  |  |  |  |  | HS |  |
|  | 16 | 0.02 |  | **HS** | HS |  |  |  |  | HS |  |  |  |  |  |
|  | 17 | **0.60** | HS | **FS** | HS |  | HS |  | HS |  |  |  |  |  |  |
|  | 18 | 0.24 | HS | **HS** |  |  |  |  |  |  |  |  |  |  |  |
|  | 19 | 0.39 |  |  |  |  | **HS** |  | HS |  |  |  |  |  |  |
|  | 20 | 0.20 |  |  | HS | HS | HS |  |  |  |  |  |  |  | HS |
|  | 23 | **0.52** |  |  | **HS** | **HS** |  |  |  |  |  |  |  |  |  |
|  | 24 | 0.41 |  |  |  | HS | **FS** |  | HS |  |  |  |  |  |  |
|  | 26 | 0.37 |  |  | **HS** |  |  | **HS** |  | **FS** |  |  |  | HS |  |
|  | 27 | 0.11 |  | **HS** | **FS** |  |  | **HS** |  |  |  |  |  |  |  |
|  | 28 | 0.27 |  |  | **FS** |  |  | **HS** |  | **FS** |  |  |  |  |  |
|  | 29 | 0.26 |  | **FS** |  | HS |  | **FS** | HS |  |  |  |  |  |  |
|  | 30 | 0.40 |  |  |  |  | HS |  | **PO** | **FS** |  |  |  |  |  |
|  | 32 | 0.43 |  |  |  |  |  | **FS** | **HS** | **PO** |  |  |  |  |  |
|  | 33 | 0.10 |  |  |  |  | **PO** | **PO** | **FS** |  |  |  |  |  |  |
|  | 34 | 0.13 |  |  |  | HS |  | **PO** | HS | **FS** |  |  | HS |  |  |
|  | 36 | 0.37 |  |  |  |  |  |  | **FS** | HS | HS | HS |  |  | HS |
|  | 38 | 0.24 |  |  |  |  |  |  |  |  |  |  |  |  |  |
|  | 39 | 0.22 |  |  | **FS** |  |  |  |  |  |  |  |  | HS |  |
|  | 40 | **0.52** |  |  |  |  | **HS** |  |  |  |  |  |  |  |  |
|  | 41 | 0.37 |  |  |  |  |  |  | HS |  | HS |  | HS |  | HS |
|  | 42 | 0.23 | **FS** |  |  | **HS** |  |  |  |  |  |  | HS |  |  |
|  | 43 | 0.41 |  | HS |  |  | **PO** |  |  | HS |  |  |  |  |  |
|  | 44 | 0.35 |  |  |  |  |  |  | HS |  |  | **HS** | HS |  |  |
|  | 45 | 0.23 |  |  |  |  |  |  |  | **FS** | **PO** | HS | HS |  | HS |
|  | 48 | **0.62** |  |  |  |  |  |  |  |  |  | HS | **FS** |  |  |
|  | 49 | 0.27 |  |  | HS |  |  |  |  |  | **HS** | **PO** |  |  |  |
|  | 50 | **1.00** |  |  |  |  |  |  |  |  | **FS** |  |  |  |  |
|  | 51 | **0.51** |  |  |  |  |  |  |  |  | HS |  |  |  |  |
|  | 52 | 0.22 |  |  |  |  |  |  |  |  |  | **FS** |  |  |  |

|  |  |  |  |  |  |  |  |  |  |  |  |  |
| --- | --- | --- | --- | --- | --- | --- | --- | --- | --- | --- | --- | --- |
| 53 | 0.36 |  |  |  |  |  |  |  |  |  |  | HS |
| 54 | 0.49 |  |  |  |  |  |  |  |  |  |  |  |
| 55 | 0.36 |  |  |  |  |  |  |  |  |  | **FS** |  |
| 56 | 0.49 |  |  |  |  |  | HS |  |  |  | HS | HS |
| 57 | 0.49 |  |  |  | HS |  |  |  |  |  | HS |  |
| 58 | 0.11 |  |  |  | HS |  | HS |  |  |  | HS | **HS** |
| 59 | 0.11 |  |  |  | **HS** | HS | **FS** |  | HS |  | HS |  |
| 61 | 0.36 |  |  |  |  | **HS** | **FS** |  |  | **HS** |  |  |
| 62 | 0.34 |  |  |  |  |  |  |  | HS | **HS** |  | **FS** |
| 63 | 0.22 |  |  | HS |  |  |  |  |  | HS |  |  |
| 64 | 0.36 |  |  |  |  |  |  |  |  | HS | **FS** | **HS** |
| 65 | 0.47 |  |  |  |  |  |  |  |  |  |  |  |
| 66 | 0.40 |  |  |  |  |  |  |  |  | **FS** | **FS** |  |
| 67 | **0.68** |  |  |  |  |  |  |  |  | HS |  | HS |
| 68 | 0.49 |  |  |  |  |  |  | HS |  |  |  |  |
| 69 | 0.49 |  |  |  |  |  |  |  |  | **FS** |  | HS |

|  |  |  |  |  |  |  |  |  |  |  |  |  |  |  |
| --- | --- | --- | --- | --- | --- | --- | --- | --- | --- | --- | --- | --- | --- | --- |
| 70 | 0.28 |  |  |  |  |  |  |  |  | HS | HS |  |  |  |
| 71 | 0.33 |  |  | HS | HS |  |  |  |  |  |  | **PO** |  | HS |
| 72 | 0.25 |  |  | **FS** |  |  |  |  |  |  |  |  | HS | HS |
| 73 | 0.22 |  |  | HS | HS |  | HS | HS |  |  |  |  | **PO** |  |
| 74 | 0.33 | HS |  |  |  |  |  | HS |  |  |  |  | **FS** |  |
| 76 | 0.22 |  | HS |  |  |  |  |  |  |  |  |  |  | **HS** |
| 77 | 0.40 |  |  |  | **FS** | HS | **HS** | HS |  |  |  |  | **HS** |  |
| 78 | 0.16 |  |  |  |  |  |  |  |  |  | HS |  |  |  |
| 79 | 0.21 |  | HS |  |  |  |  |  |  |  |  |  |  | **FS** |
| 80 | 0.45 |  |  |  |  | HS | **HS** |  |  |  | HS |  |  |  |
| 81 | 0.14 | HS |  |  |  |  |  |  |  | HS |  |  |  | HS |
| 82 | 0.12 |  |  |  |  |  |  |  |  |  |  |  |  | **FS** |
| 84 | 0.39 |  |  |  |  | HS |  |  |  |  |  |  |  | **FS** |
| 85 | 0.41 | **HS** | HS |  |  |  |  |  |  |  |  |  |  |  |
| 86 | 0.41 |  | **HS** |  |  |  |  |  |  |  |  |  |  | **FS** |

Only three kinds of relationship are indicated: Half-sibling (HS), full-sibling (FS), and parent-offspring (PO) meaning that when a pair has no indication it is unrelated. Pedigree indicated in bold corresponds to result of highest likelihood prediction while the grey color
indicates that this pedigree changed to unrelated after we applied the likelihood ratio test. Females highlighted in grey have no kin relationship with the 13 males. For all individuals, homozygosity (*HL*) is given as informative and values above 0.5 are indicated in bold.

## Discussion

There are numerous studies on *Trachemys* spp, including topics such as general ecology [25, 55–56], biogeography [57], systematics [17, 58], and even as an invasive species [59–61]. This is probably because many species in this genus are included within the IUCN Red List and TFTSG list (Tortoise and Freshwater Turtle Specialist Group) or are not yet evaluated [62]. Genetic studies are also largely represented, generally focusing on phylogenic or biogeographic considerations [13–14, 16], for hybridization process [15], or even identifying genetic damage under radioactive effect [63]; however, population genetic studies are rarer [64–67]. Particularly, for *Trachemys venusta*, no genetic population study is reported to date. We present, the first results related to the population genetics of wild population of *T. venusta venusta* in Mexico compared to founder individuals of a captive group (Mexican UMA).

With the exception of some studies that present low genetic diversity (value of *H*_e_ < 0.4; [68–69]), freshwater turtles exhibit high values of genetic diversity (generally *H*_e_ > 0.6-0.7; see Table 4 in [69] for examples, and [70]) based on microsatellites. Our values could be considered lower (*H*_e_ < 0.6) than many other freshwater turtles or even threatened species (see Table 4 in [69] for examples, and [70–71]), and are in agreement with values for wild populations of *Elusor macrurus* Cann & Legler 1994 (Testudines, Chelidae) an endangered Australian freshwater turtle [72]. The sole genetic study that reports value of genetic diversity for *Trachemys* spp. based on microsatellites showed low values for *T. taylori* Legler, 1960 (*H*_e_ from 0.442 to 0.569; *H*_o_ from 0.458 to 0.590) and very high values for *T. scripta elegans* Wied-Neuwied, 1839 (*H*_e_ = 0.809; *H*_o_ = 0.828) [67]. Other population genetic studies that report levels of genetic diversity for *Trachemys* spp used allozymes which makes comparison with microsatellites data difficult. In this respect, all studies showed very low values of genetic diversity (*H* < 0.15; [64–66]). The relatively low value of genetic diversity observed for wild population of *T. venusta venusta* compared to other turtle species, suggests that this species deserves reinforced attention and probably higher level of protection considering the threshold of 0.54 proposed by Willoughby et al. [73] as a value to consider a species as Critically Endangered [72].

Values of genetic diversity for captive population were slightly higher than wild population as reported for *Elusor macrurus* [72] and for *Dermatemys mawii* Gray, 1847 (Testudines, Dermatemydidae) [74]. Because founder individuals of the UMAs (captive population) come from two distinct localities: Pomposu that is within the Grijalva River Sub-Basin and El Espino in the Usumacinta River Sub-Basin (see Fig 1), the genetic pool of those founders could be higher due to the mix of genetic sources. This situation was reported for *Lithobates sevosus* Goin & Netting, 1940 (Anura, Ranudae), a critically endangered frog from the southeastern USA [75], and for *Dermatemys mawii* a critically endangered freshwater turtle [74], suggesting that the captive population is genetically representative of natural populations [75] and could be used to found new wild populations [76]. Also, founder individuals are approximately 40 years of age, and genetic diversity represents the situation from several decades ago. The genetic diversity of the wild population of *T. venusta venusta* may have declined since the UMA was founded, probably due to habitat fragmentation and water pollution [19]. Indeed, the state of Tabasco lost around 60% of its wetlands at the beginning of the 21st century, mainly due to anthropogenic activities such as the oil industry, the establishment of new crop areas and grassland for livestock use, road construction, population growth, and increased pollution derived from these activities [20, 77]. Additionally, in Tabasco state, freshwater turtles are subject to intense hunting and illegal trading and trafficking [78–80] that has contributed to a decline in population numbers leading to a loss of genetic diversity. It is reported that in Villahermosa and surrounding cities (e.g., Nacajuca, Comalcalco, Jalpa de Méndez, and Cunduacán) freshwater turtles were abundant, but today it is difficult to find turtles such as *T. venusta* [80]. Many species of turtles are characterized by a long generation time (often > 25 years), making a loss of genetic diversity or genetic differentiation among populations due to recent and/or anthropogenic events (e.g., habitat fragmentation) very difficult or impossible to detect (see examples in [70]). However, sexual maturity of *T. venusta* has been evaluated at approximately four to seven years [81] which results in five to 10 generations (40 years) between UMA founder individuals and the wild population analyzed in this study, thus allowing the observation of a loss of genetic diversity in the wild population due to recent anthropogenic pressure.

Although captive population (UMA) did not demonstrate a significant loss of heterozygotes, wild population shows significant loss of heterozygotes that could reveal high level of inbreeding and small population size that are probably a consequence of recent anthropogenic pressures (see above) on freshwater turtles in Tabasco state. Several species of freshwater turtles, such as *Apalone spinifera emoryi* Agassiz, 1857 (Testudines, Trionychidae), *Mesoclemmys dahli* Zangerl & Medem, 1958 (Testudines, Chelidae), *Chrysemys p. picta* Schneider, 1783 (Testudines, Emydidae), and *Clemmys guttata* Schneider, 1792 (Testudines, Emydidae) have shown evidence of genetic isolation, genetic differentiation as well as modest to high inbreeding rates, but surprisingly the values of heterozygosity in these species are medium to high (0.6-0.7) despite experiencing anthropogenic pressures [82–84]. As mentioned before, the decrease in genetic diversity of many species of turtles, even after a prolonged decrease in population size, may not be observed due to their long generation times and late maturity associated with chelonian life history [84–86].

Bayesian and PCoA analysis of genetic structure among wild and captive populations revealed a relatively good separation between both probably reflecting the low-connectivity among sites where founder individuals originated. Indeed, as mentioned before, the Pomposu locality depends on the Grijavlva River Sub-Basin as well as wild populations considered in this work (captive individuals showing a dominant wild genetic profile in analysis). El Espino depends on the Usumacinta River Sub-Bassin (see Fig 1) and could be represented by captive individuals with the other genetic profile in Structure analysis.

### Kindship analysis

The relationship coefficient evaluated for captive population (Tabasco State Government UMA) suggest a relatively low level of kinship (considering all pairs: female-male, female-female, and male-male) which probably reflects that the founding individuals of the UMA came from different populations. However, we highlighted the differences observed between programs (Storm and ML-Relate) and between methods used in ML-Relate (using or not the likelihood ratio test to evaluate the more probable relationship) which can give values of single to double (e.g., 45% for unrelated using Storm and 91% using ML-Relate with ratio test).

From all founder females of the UMA (n = 73), the stricter analysis provided by ML-Related enabled the identification of 19 females that present no kin relationship with males (n = 13). This could result in a total of 247 potential dyads to establish an optimal breeding program in the UMA. In order to develop the optimal reproduction in a captive population, the number and the identity of the founders is a key factor for the genetic pool [87]. According to these authors, only 15 individuals are necessary to preserve a *H_e_* of 0.54 based on a revision of 188 studies where genetic conservation approaches are used for breeding programs. Considering this, if only unrelated founders were considered for reproduction, the captive population of Tabasco would have an optimal number of founding individuals to ensure genetic diversity. The captive population of Tabasco could be used for the reintroduction of *T. venusta venusta*, considering that this species is in decline in its natural habitat with low genetic diversity (see above). For this, we recommended the use of the optimal females (no kin relationship with all males of the UMA), and the introduction of new founders to avoid generic erosion [87]. To date, very few studies have used genetic information of founders to optimize breeding program and avoid inbreeding in turtle programs. For example, Miller et al. [2] evaluated the pedigree on three seasons of a captive breeding program of the Galápagos giant tortoise showing that the genetic diversity of the progeny was reduced in a single generation and there was a tendency towards the reduction of the physical shape when more related individuals were bred. Also, Spitzweg et al. [88] analyzed the genetic diversity and kin relationship of a turtle (*Batagur baska* Gray, 1830; Testudines, Geoemydidae) in captivity, and showed that most of the founder individuals that came from wildlife had a high kinship relation.

Successful management and conservation programs for long-lived organisms are those that recognize the need for protection and biological knowledge of all life stages of the species they breed [89]. For this reason, our results have important implications for the conservation and management of *T. venusta venusta*, as they contribute to a better selection of pairs, decreasing the possibility of inbreeding in the generations born in captivity. Furthermore, through good genetic management, individuals could be released into the wild to improve the genetic diversity of the *trivittata* Duméril & Bibron, 1835 (Testudines, Geoemydidae) is an example of a successful genetic management program, where the breeding, reproduction, and release of captive individuals has resulted in an improvement in the wild population. A captive management program has been established for this species, and it was thought that only 12 breeding turtles existed in the wildlife. Since 2002, more than 700 individuals of this species have been obtained and after selecting individuals with a high genetic diversity and reintroducing them to their natural habitat, the program reported an increase in the fertility of the eggs [90]. Furthermore, our results could be used to create predictive models of exploitation, since *T. venusta venusta* is a highly valued species for human consumption and as pet, as has been proposed for the turtle *Trachemys scripta elegans* Wied-Neuwied, 1839 (Testudines, Emydidae), that is principally exploited as a pet [91].

Finally, we believe that it is convenient to analyze the offspring of the founders already in the UMA in order to make a better use of this captive population. Another recommendation to consider in the implementation of a genetic management program for *T. venusta* could be the storage of sperm and the study of the multiple paternity, since it has been reported in several species of turtles [76] as a reproductive strategy that serves to maximize the genetic diversity of the offspring of long-lived organisms [92].

## Acknowledgments

The authors would like to thank the Mexican Commission for the Knowledge and Use of Biodiversity (CONABIO) for the funding with number NE017-project: “Establishment of a genetic management program for the species *Dermatemys mawii* (white turtle) and *Trachemys venusta* (hicotea) in Wildlife Management Units (UMA) to increase the genetic flow and connectivity of the Mesoamerican Biological Corridor in Tabasco” which allowed to carry out the present study. Thanks to the Mexican National Council of Science and Technology (CONACyT) for the maintenance grant of the Biol. Elsi Beatriz Recino Reyes for the master’s studies in Environmental Sciences at Universidad Juárez Autónoma de Tabasco, Division of Biological Sciences. The capture of organisms was done with the permit number SPGA/DGVS/01085/16 issued by the Mexican Ministry of Environment and Natural Resources (SEMARNAT). We would also like to thank Dr Humberto Hernández Trejo PhD for his contribution of information to strengthen the discussion. Thanks to Dr. Leon David for helping to produce the map of Fig 1.

## Data Accessibility Statement

The morphometric and weight data taken from *Trachemys venusta venusta* can be found at Knowledge Network for Biocomplexity Digital Repository: doi:10.5063/F1C53J44.

## Supporting information caption

**S1 Table. Analysis of molecular variance (AMOVA) for *Trachemys venusta venusta* in south Mexico**. When two groups are considered (UMA *vs* wild populations) (A), and when localities of wild individuals are included in the analysis (B).

**S1 Fig. Distribution of frequencies of internal relatedness (*IR*) for *Trachemys venusta venusta* in south Mexico**. In UMA (purple) and in wild population (blue).

**S2 Fig. Distribution of frequencies of homozygosity by loci (*HL*)** for *Trachemys venusta venusta* in south Mexico. UMA (purple) and wild population (blue).

S3 Fig. Identification of the optimal number of groups (clusters) using the Delta K method implemented in STRUCTURE HARVESTER website.

## References

1. Valiente-Banuet A, Aizen MA, Alcántara JM, Arroyo J, Cocucci A, Galetti M, et al. Beyond species loss: the extinction of ecological interactions in a changing world. Funct Ecol. 2015; 29: 299–307.

2. Miller JM, Quinzin MC, Scheibe EH, Ciofi C, Villalva F, Tapia W, et al. Genetic pedigree analysis of the pilot breeding program for the rediscovered Galapagos giant tortoise from Floreana island. J Hered. 2018; 109: 620–630.

3. Ballou JD, Lees C, Faust LJ, Long S, Lynch C, Bingaman Lackey L, et al. Demographic and genetic management of captive populations. In: Kleiman DG, Thomson KV, Keirk-Baer C, editors. Wild mammals in captivity: principles and techniques for zoo management. University of Chicago Press; 2010. pp. 219–252.

4. Ortega-Argueta A, González-Zamora A, Contreras-Hernández A. A framework and indicators for evaluating policies for conservation and development: The case of wildlife management units in Mexico. Environ Sci Policy. 2016; 63:91–100.

5. Weber M, García-Marmolejo G, Reyna-Hurtado R. The tragedy of the commons: wildlife management units in southeastern Mexico. Wildl Soc Bull. 2006; 34: 1480–1488.

6. Valdez R, Guzmán-Aranda JG, Abarca FJ, Tarango-Arámbula LA, Clemente-Sánchez F. Wildlife Conservation and Management in Mexico. Wildl Soc Bull. 2006; 34(2): 270–282.

7. SEMARNAT. Programa de Conservación de la Vida Silvestre y Diversificación Productiva en el Sector Rural. México. Secretaria de Medio Ambiente, Recursos Naturales y Pesca, México; 1997.

8. de Benito RR. Las unidades de manejo para la conservación de vida silvestre y el Corredor Biológico Mesoamericano México. Comisión Nacional para el Conocimiento y Uso de la Biodiversidad. México; 2009.

9. Ballou JD, Lacy RC. Identifying genetically important individuals for management of genetic variation in pedigreed populations. In: Ballou JD, Gilpin M, Foose T, editors. Population management for survival and recovery. Columbia University Press; 1995. pp. 76–111.

10. Williams SE, Hoffman EA. Minimizing genetic adaptation in captive breeding programs: a review. Biol Conserv. 2009; 142: 2388–2400.

11. Zarza E, Reynoso VH, Emerson BC. Genetic tools for assisting sustainable management and conservation of the spiny-tailed iguana, *Ctenosaura pectinata*. Herpetol Conserv Biol. 2016; 11: 255–264.

12. Seidel ME. Taxonomic observations on extant species and subspecies of slider turtles, genus *Trachemys*. J Herpetol. 2002; 36: 285–292.

13. Parham JF, Papenfuss TJ, Buskirk JR, Parra-Olea G, Chen JY, Simison WB. *Trachemys ornata* or not *ornata*: reassessment of a taxonomic revision for Mexican *Trachemys*. Proc Calif Acad Sci. 2015; 62: 359–367.

14. Fritz U, Stuckas H, Vargas-Ramírez M, Hundsdörfer AK, Maran J, Päckert M. Molecular phylogeny of Central and South American slider turtles: implications for biogeography and systematics (Testudines: Emydidae: *Trachemys*). J Zool Syst Evol Res. 2012; 50: 125–136.

15. Parham JF, Papenfuss TJ, Van Dijk PP, Wilson BS, Marte C, Schettino LR, et al. Genetic introgression and hybridization in Antillean freshwater turtles (*Trachemys*) revealed by coalescent analyses of mitochondrial and cloned nuclear markers. Mol Phylogenet Evol. 2013; 67: 176–187.

16. Vargas-Ramírez M, del Valle C, Ceballos CP, Fritz U. *Trachemys medemi* n. sp. from northwestern Colombia turns the biogeography of South American slider turtles upside down. J Zool Syst Evol Res. 2017; 55: 326–339.

17. Ernst CH, Seidel ME. Trachemys venusta. In: Price AH, editor. Catalogue of American Amphibians and Reptiles (CAAR). Society for the Study of Amphibian and Reptiles; 2006. pp. 832.1-832.12

18. Guevara Chumacero M, Pichardo Fragoso A, Martínez Cornelio M. La tortuga en Tabasco: comida, identidad y representación. Estudios de Cultura Maya. 2017; 49: 97–122.

19. Tudela F. La modernización forzada del trópico: El caso de Tabasco, proyecto integrado del Golfo. IFIAS. México, D.F; 1992

20. Palomeque de la Cruz MA, Galindo Alcántara A, Sánchez AJ, Escalona Maurice MJ. Pérdida de humedales y vegetación por urbanización en la cuenca del río Grijalva, México. Investigaciones Geográficas. 2017; 68: 151–172.

21. Flores-Villela O, Canseco-Márquez L. Nuevas especies y cambios taxonómicos para la herpetofauna de México. Acta Zool Mex (n.s.). 2004; 20: 115–144.

22. SEMARNAT. Secretaría de Medio Ambiente y Recursos Naturales. Norma Oficial Mexicana NOM-059-SEMARNAT-2010. Diario Oficial de la Federación (DOF); 2010.

23. Blouin MS. DNA-based methods for pedigree reconstruction and kinship analysis in natural populations. Trends Ecol Evol. 2003; 18: 503–511.

24. Vogt RC. New methods for trapping aquatic turtles. Copeia. 1980; 1980: 368–371.

25. Gibbons JW. Life history and ecology of the slider turtle. Washington, DC: Smithsonian Institution Press; 1990.

26. SEMARNAT. Memorias del Taller de capacitación para la conservación y aprovechamiento sustentable de tortugas dulceacuícolas del sur-sureste de México. Catemaco, Veracruz; 2009.

27. Proebstel BS, Evas RP, Shiozawa DK, Williams RN. (1993). Preservation of nonfrozen tissue samples from a salmonine fish *Brachymystax lenok* (Pallas) for DNA analysis. J Ichthyol. 1993; 32: 9–17.

28. Simison WB, Sellas AB, Feldheim KA, Parham JF. Isolation and characterization of microsatellite markers for identifying hybridization and genetic pollution associated with red-eared slider turtles (*Trachemys scripta elegans*). Conserv Genet Res. 2013; 5: 1139–1140.

29. Rozen S, Skaletsky HJ. Primer3 on the WWW for general users and for biologist programmers. In: Krawetz S, Misener S, editors. Bioinformatics methods and protocols: methods in molecular biology. Humana Press, Totowa; 2000. pp. 365–386.

30. Valière N. GIMLET: a computer program for analysing genetic individual identification data. Mol Ecol Notes. 2002; 2: 377–379.

31. Bonin A, Bellemain E, Bronken Eidesen P, Pompanon F, Brochmann C, et al. How to track and assess genotyping errors in population genetics studies. Mol Ecol. 2004; 13: 3261–3273.

32. Machkour-M’Rabet S, Cruz-Medina J, García-De León FJ, De Jesús-Navarrete A, Hénaut Y. Connectivity and genetic structure of the queen conch on the Mesoamerican Reef. Coral Reefs. 2017; 36: 535–548.

33. Peakall R, Smouse PE. GenAlEx 6.5: genetic analysis in Excel. Population genetic software for teaching and research-an update. Bioinformatics. 2012; 28: 2537–2539.

34. Hammer O, Harper DAT, Ryan PD. PAST: Paleontological Statistics Software Package for Education and Data Analysis. Palaeontol Electron. 2001; 4: 9pp.

35. Kalinowski ST. Counting alleles with rarefaction: private alleles and hierarchical sampling designs. Conserv Genet. 2004; 5: 539–543.

36. Kalinowski ST. Hp-rare 1.0: a computer program for performing rarefaction on measures of allelic richness. Mol Ecol Notes. 2005; 5: 187–189.

37. Kalinowski ST, Taper ML, Marshall TC. Revising how the computer program CERVUS accommodates genotyping error increases success in paternity assignment. Mol Ecol. 2007; 16: 1099–1106.

38. Amos W, Wilmer JW, Fullard K, Burg TM, Croxall JP, Bloch D, et al. The influence of parental relatedness on reproductive success. Proc R Soc Lond B: Biol Sci. 2001; 268: 2021–2027.

39. Aparicio JM, Ortego J, Cordero PJ. What should we weigh to estimate heterozygosity, alleles or loci? Mol Ecol. 2006; 15: 4659–4665.

40. Raymond M, Rousset, F. GENEPOP (version 1.2): population genetics software for exact test and ecumenicism. J Hered. 1995; 86: 248–249.

41. Rousset, F. Genepop’007: a complete reimplementation of the Genepop software for Windows and Linux. Mol Ecol Res. 2008; 8: 103–106.

42. Chapuis MP, Estoup A. Microsatellite null alleles and estimation of population differentiation. Mol Biol Evol. 2006; 24: 621–631.

43. Do C, Waples RS, Peel D, Macbeth GM, Tillett BJ, Ovenden JR. NeEstimator v2: re-implementation of software for the estimation of contemporary effective population size (Ne) from genetic data. Mol Ecol Res. 2014; 14: 209–214.

44. Waples RS, Do CHI. LDNE: a program for estimating effective population size from data on linkage disequilibrium. Mol Ecol Res. 2008; 8: 753–756.

45. Cornuet JM, Luikart G. Description and power analysis of two tests for detecting recent population bottlenecks from allele frequency data. Genetics. 1996; 144: 2001–2014.

46. Piry S, Luikart G, Cornuet JM. BOTTLENECK: a computer program for detecting recent reductions in the effective population size using allele frequency data. J Hered. 1999; 90: 502– 503.

47. Pritchard JK, Stephens M, Donnelly P. Inference of population structure using multilocus genotype data. Genetics. 2000; 155: 945–959.

48. Evanno G, Regnaut S, Goudet J. Detecting the number of clusters of individuals using the software STRUCTURE: a simulation study. Mol Ecol. 2005; 14: 2611–2620.

49. Earl DA, Von Holdt BM. STRUCTURE HARVESTER: A website and program for visualizing STRUCTURE output and implementing the Evanno method. Conserv Genet Res. 2011; 4: 359–361.

50. Frasier TR. STORM: software for testing hypotheses of relatedness and mating patterns. Mol Ecol Res. 2008; 8: 1263–1266.

51. Valiere N, Bonenfant C, Toïgo C, Luikart G, Gaillard JM, Klein F. Importance of a pilot study for non-invasive genetic sampling: genotyping errors and population size estimation in red deer. Conserv Genet. 2007; 8: 69–78.

52. Říčanová Š, Bryja J, Cosson JF, Gedeon C, Choleva L, Ambros M, et al. Depleted genetic variation of the European ground squirrel in Central Europe in both microsatellites and the major histocompatibility complex gene: implications for conservation. Conserv Genet. 2011; 12: 1115–1129.

53. Queller DC, Goodnight KF. Estimating relatedness using genetic markers. Evolution. 1989; 43: 258–275.

54. Kamel SJ, Hughes AR, Grosberg RK, Stachowicz JJ. Fine-scale genetic structure and relatedness in the eelgrass Zostera marina. Mar Ecol Prog Ser. 2012; 447: 127–137.

55. Gradela A, Santiago TOC, Pires IC, Silva ADCS, de Souza LC, de Faria MD, et al. Sexual Dimorphism in Red-Eared Sliders (Trachemys scripta elegans) from the Wild Animal Triage Center of the Tiete Ecological Park, São Paulo, Brazil. Acta Sci Vet. 2017; 45: 1-10.

56. Works AJ, Olson DH. Diets of Two Nonnative Freshwater Turtle Species (*Trachemys scripta* and *Pelodiscus sinensis*) in Kawai Nui Marsh, Hawaii. J Herpetol. 2018; 52: 444–452.

57. Ennen JR, Matamoros WA, Agha M, Lovich JE, Sweat SC, Hoagstrom CW. Hierarchical, quantitative biogeographic provinces for all North American turtles and their contribution to the biogeography of turtles and the continent. Herpetol Monogr. 2017; 31: 142–168.

58. McCord WP, Joseph-Ouni M, Hagen C, Blanck T. Three new subspecies of *Trachemys venusta* (Testudines: Emydidae) from Honduras, northern Yucatán (Mexico), and pacific coastal Panama. Reptilia. 2010; 71: 39–49.

59. Cadi A, Joly P. Competition for basking places between the endangered European pond turtle (*Emys orbicularis galloitalica*) and the introduced red-eared slider (*Trachemys scripta elegans*). Can J Zool. 2003; 81: 1392–1398.

60. Rodrigues JFM, Coelho MTP, Diniz-Filho JAF. Exploring intraspecific climatic niche conservatism to better understand species invasion: the case of Trachemys dorbigni (Testudines, Emydidae). Hydrobiologia. 2016; 779: 127–134.

61. Martins BH, Azevedo F, Teixeira J. First reproduction report of *Trachemys scripta* in Portugal-Ria Formosa Natural Park, Algarve. Limnetica. 2018; 37: 61–67.

62. Rhodin AG, Stanford CB, Van Dijk PP, Eisemberg C, Luiselli L, Mittermeier RA, et al. Global conservation status of turtles and tortoises (Order Testudines). Chelonian Conserv Bi. 2018; 17: 135–161.

63. Lamb T, Bickham JW, Gibbons JW, Smolen MJ, McDowell S. Genetic damage in a population of slider turtles (*Trachemys scripta*) inhabiting a radioactive reservoir. Arch Environ Con Tox. 1991; 20: 138–142.

64. Scribner KT, Smith MH, Gibbons JW. Genetic differentiation among local populations of the yellow-bellied slider turtle (*Pseudemys scripta*). Herpetologica. 1984; 40: 382–387.

65. Smith MH, Scribner KT. Population genetics of the slider turtle. In Gibbons JW, editor. Life History and Ecology of the Slider Turtle. Smithsonian Institution Press, Washington, D.C.; 1990. pp. 74-81.

66. Martínez LM, Bock BC, Páez VP. Population genetics of the slider turtle (*Trachemys scripta callirostris*) in the Mompos Depression, Colombia. Copeia. 2007; 2007: 1001–1005.

67. McGaugh SE. Comparative population genetics of aquatic turtles in the desert. Conserv Genet. 2012; 13: 1561–1576.

68. Echelle AA, Hackler JC, Lack JB, Ballard SR, Roman J, Fox SF, et al. Conservation genetics of the alligator snapping turtle: cytonuclear evidence of range-wide bottleneck effects and unusually pronounced geographic structure. Conserv Genet. 2010; 11: 1375–1387.

69. Vargas-Ramírez M, Stuckas H, Castano-Mora OV, Fritz U. Extremely low genetic diversity and weak population differentiation in the endangered Colombian river turtle *Podocnemis lewyana* (Testudines: Podocnemididae). Conserv Genet. 2012; 13: 65–77.

70. Davy CM, Bernardo PH, Murphy RW. A Bayesian approach to conservation genetics of Blanding’s turtle (*Emys blandingii*) in Ontario, Canada. Conserv Genet. 2014; 15: 319–330.

71. Petre C, Selman W, Kreiser B, Pearson SH, Wiebe JJ. Population genetics of the diamondback terrapin, *Malaclemys terrapin*, in Louisiana. Conserv Genet. 2015; 16: 1243–1252.

72. Schmidt DJ, Espinoza T, Connell M, Hughes JM. Conservation genetics of the Mary River turtle (*Elusor macrurus*) in natural and captive populations. Aquat Conserv 2018; 28: 115–123.

73. Willoughby JR, Sundaram M, Wijayawardena BK, Kimble SJ, Ji Y, Fernandez NB, et al. The reduction of genetic diversity in threatened vertebrates and new recommendations regarding IUCN conservation rankings. Biol Conserv. 2015; 191: 495–503.

74. Gallardo-Alvárez MI, Lesher-Gordillo JM, Machkour-M’Rabet S, Zenteno-Ruiz CE, Olivera-Gómez LD, del Rosario Barragán-Vázquez M, et al. Genetic diversity and population structure of founders from wildlife conservation management units and wild populations of critically endangered *Dermatemys mawii*. Glob Ecol Conserv. 2019; 19: e00616.

75. Hinkson KM, Henry NL, Hensley NM, Richter SC. Initial founders of captive populations are genetically representative of natural populations in critically endangered dusky gopher frogs, *Lithobates sevosus*. Zoo Biol. 2016; 35: 378–384.

76. Edwards T, Cox EC, Buzzard V, Wiese C, Hillard LS, Murphy RW. Genetic assessments and parentage analysis of captive Bolson tortoises (*Gopherus flavomarginatus*) inform their “rewilding” in New Mexico. PLOS ONE. 2014; 9: e102787.

77. Melchor H, Isela G, Ruíz Rosado O, Sol Sánchez Á, Valdez Hernández JI. Cambios de uso del suelo en manglares de la costa de Tabasco. Rev Mexicana cienc agríc. 2016; 7(SPE14): 2757–2767.

78. PROFEPA. Asegura PROFEPA 30 tortugas en Tabasco. PROFEPA.gob. 2014. Available from: http://www.profepa.gob.mx/innovaportal/v/5687/1/mx.wap/asegura_profepa_30_tortugas_en_tabasco.html

79. PROFEPA. (2015). Remite PROFEPA a MPF a sujeto en posesión de ejemplares de vida silvestre en situación de riesgo. PROFEPA.gob. 2015. Available from: http://www.profepa.gob.mx/innovaportal/v/7159/1/mx/remite_profepa_a_mpf_a_sujeto_en_posesion_de_ejemplares_de_vida_silvestre_en_situacion_de_riesgo.html

80. Reynoso VH, Vázquez Cruz ML, Rivera Arroyo RC. Estado de conservación, uso, gestión, comercio y cumplimiento de los criterios de inclusión a los Apéndices de la CITES para las especies *Claudius angustatus* y *Staurotypus triporcatus*. Universidad Nacional Autónoma de México. Instituto de Biología. Informe final SNIBCONABIO, Ciudad de México. 2016. Available from: http://www.conabio.gob.mx/institucion/proyectos/resultados/InfMM009.pdf

81. Torres KL, Franyutti AAH, Aranzábal MDCU, Vidal UH. Características reproductoras de la tortuga dulceacuícola hicotea (*Trachemys venusta*). Kuxulkab’. 2011; 17: 43–49.

82. Mali I, Villamizar-Gomez A, Guerra TM, Vandewege MW, Forstner MR. Population genetics of Texas spiny softshell turtles (*Apalone spinifera emoryi*) under various anthropogenic pressures in two distinct regions of their range in Texas. Chelonian Conserv Bi. 2015a; 14: 148–156.

83. Gallego-García N, Vargas-Ramírez M, Forero-Medina G, Caballero S. Genetic evidence of fragmented populations and inbreeding in the Colombian endemic Dahl’s toad-headed turtle (*Mesoclemmys dahli*). Conserv Genet. 2018; 19: 221–233.

84. Buchanan SW, Kolbe JJ, Wegener JE, Atutubo JR, Karraker NE. A Comparison of the Population Genetic Structure and Diversity between a Common (*Chrysemys p. picta*) and an Endangered (*Clemmys guttata*) Freshwater Turtle. Diversity. 2019; 11: 99.

85. Kuo CH, Janzen FJ. Genetic effects of a persistent bottleneck on a natural population of ornate box turtles (*Terrapene ornata*). Conserv Genet. 20047; 5: 425–437.

86. Willoughby JR, Sundaram M, Lewis TL, Swanson BJ. Population decline in a long-lived species: the wood turtle in Michigan. Herpetologica. 2013; 69: 186–198.

87. Witzenberger KA, Hochkirch A. Ex situ conservation genetics: a review of molecular studies on the genetic consequences of captive breeding programmes for endangered animal species. Biodivers Conserv. 2011; 20: 1843–1861.

88. Spitzweg C, Praschag P, DiRuzzo S, Fritz U. Conservation genetics of the northern river terrapin (*Batagur baska*) breeding project using a microsatellite marker system. SALAMANDRA. 2018; 54: 63–70.

89. Congdon JD, Dunham AE, van Loben Sels RC. Delayed sexual maturity and demographics of Blanding’s turtles (*Emydoidea blandingii*): implications for conservation and management of long-lived organisms. Conserv Biol. 1993; 7: 826–833.

90. Çilingir FG, Rheindt FE, Garg KM, Platt K, Platt SG, Bickford DP. Conservation genomics of the endangered Burmese roofed turtle. Conserv Biol. 2017; 31: 1469–1476.

91. Mali I, Wang HH, Grant WE, Feldman M, Forstner MR. Modeling commercial freshwater turtle production on US farms for pet and meat markets. PLOS ONE. 2015b; 10: e0139053.

92. Davy CM, Edwards T, Lathrop A, Bratton M, Hagan M, Henen B, et al. Polyandry and multiple paternities in the threatened Agassiz’s desert tortoise, *Gopherus agassizii*. Conserv Genet. 2011; 12: 1313.

